# A nuclear cAMP microdomain suppresses tumor growth by Hippo pathway inactivation

**DOI:** 10.1101/2021.11.15.468656

**Authors:** Marek M. Drozdz, Ashley S. Doane, Rached Alkallas, Garrett Desman, Rohan Bareja, Michael Reilly, Jakyung Bang, Maftuna Yusupova, Jaewon You, Jenny Z. Wang, Akansha Verma, Kelsey Aguirre, Elsbeth Kane, Ian R. Watson, Olivier Elemento, Elena Piskounova, Taha Merghoub, Jonathan H. Zippin

## Abstract

cAMP signaling pathways are critical for both oncogenesis and tumor suppression. cAMP signaling is localized to multiple spatially distinct microdomains, but the role of cAMP microdomains in cancer cell biology is poorly understood. We developed a tunable genetic system that allows us to activate cAMP signaling in specific microdomains. We uncovered a previously unappreciated nuclear cAMP microdomain that functionally activates a tumor suppressive pathway in a broad range of cancers by inhibiting YAP, a key effector protein of the Hippo pathway, inside the nucleus. We show that nuclear cAMP induces a LATS-dependent pathway leading to phosphorylation of nuclear YAP solely at serine 397, export of YAP from the nucleus, without YAP protein degradation. Thus, nuclear cAMP inhibition of nuclear YAP is distinct from other known mechanisms of Hippo regulation. Pharmacologic targeting of specific cAMP microdomains remains an untapped therapeutic approach for cancer, and since Hippo pathway deregulation can lead to oncogenesis and chemotherapeutic resistance, drugs directed at the nuclear cAMP microdomain may provide new avenues for the treatment of cancer.

## Introduction

Cyclic AMP (cAMP), a ubiquitous second messenger, can induce a wide-range of cellular responses including proliferation, differentiation, and migration in response to both extracellular (e.g., hormone) or intracellular (e.g., pH) signals(Chang and Oude-Elferink, 2014; Rehmann et al., 2007; Sassone-Corsi, 2012). cAMP is known to induce both tumor promoting and tumor suppressive effects in a variety of cancers(Baljinnyam et al., 2010; Coles et al., 2020; Johannessen et al., 2013; Kloster et al., 2008; Lyons et al., 2013; Michalides et al., 2004; Patra et al., 2018; Sheppard et al., 1984), also reviewed here(Fajardo et al., 2014). How these seemingly opposing roles of cAMP exist in cancer cells has remained unresolved. It has been proposed that these differential effects of cAMP in cancer could be explained by the existence of distinct microdomains of cAMP signaling, each having disparate effects on tumor cell biology(Beavo and Brunton, 2002; Desman et al., 2014; Musheshe et al., 2018; Torres-Quesada et al., 2017; Zaccolo, 2011).

In mammalian cells, two classes of adenylyl cyclases synthesize cAMP: transmembrane adenylyl cyclases (tmAC), which generate cAMP exclusively at the plasma membrane and endosomes, and the soluble adenylyl cyclase (sAC), which generates cAMP within both the cytoplasm and organelles, specifically the mitochondria and the nucleus(Cooper and Crossthwaite, 2006; Tresguerres et al., 2011). Adjacent to each source of cAMP are phosphodiesterases (PDEs), which prevent the diffusion of cAMP from one microdomain to another(Lohse et al., 2017; Musheshe et al., 2018). Previous studies investigating the role of cAMP in tumor cell biology have focused on the role of only the tmAC class of enzymes and PDEs using reagents that can affect multiple cAMP microdomains (e.g., cAMP analogs, PDE inhibitors, and forskolin), which can lead to the loss of cAMP signaling specificity(Halls and Cooper, 2017; Zaccolo and Pozzan, 2002). Furthermore, multiple reports have shown that effector proteins of cAMP signaling (e.g., protein kinase A) can be spatially restricted even in cancer(Smith et al., 2017; Zhang et al., 2020a) suggesting that activation of different sources of cAMP in cancer cells may play an important regulatory role. However, the role of individual microdomain sources of cAMP, especially those that reside inside organelles, in tumor cell biology has remained poorly understood up to date, probably due to the lack of appropriate means to study individual sources of cAMP in distinct microdomains.

The Hippo pathway plays an important regulatory role in cancer cell proliferation, invasion, and apoptosis(Yu et al., 2015a). The Hippo signaling cascade is induced by a plethora of upstream chemical and physical stimuli and controls cancer cell proliferation and invasion, principally via the activity of YAP and TAZ(Zanconato et al., 2019), the two main transcriptional coactivators of the Hippo pathway. It is reported that cAMP generated by tmACs in the cytoplasm can lead to protein kinase A (PKA)-dependent phosphorylation of LATS and subsequent phosphorylation and inactivation of YAP(Kim et al., 2013; Yu et al., 2013). However, even though the main site of YAP and TAZ function is inside the nucleus, how YAP or TAZ is regulated within the nucleus remains poorly described. Interestingly, a nuclear localized LATS kinase is reported(Li et al., 2014), but whether cAMP can regulate LATS in the nucleus is not defined. A nuclear cAMP signaling microdomain consisting of sAC and PKA is known(Sample et al., 2012a; Zippin et al., 2004b) suggesting that this cAMP signaling cascade might influence nuclear YAP or TAZ activity via LATS.

Here we examined the effects of multiple distinct cAMP microdomains on cancer cell growth to address how this single second messenger can lead to such disparate effects in cancer. To address this fundamental question, we developed a unique tunable genetic system that allows for the activation of spatially and temporally distinct cAMP microdomain signaling within a cell and in mice. We discovered the existence of a tumor suppressive pathway evoked in multiple cancers solely by the sAC nuclear cAMP microdomain. We demonstrated that nuclear cAMP signaling induces the nuclear PKA-dependent, LATS-dependent phosphorylation of nuclear YAP solely at serine 397 (S397) without phosphorylation of YAP at serine 127 (S127) or phosphorylation of TAZ. Phosphorylation of nuclear YAP at S397 induced YAP export from the nucleus without affecting protein stability. Phosphorylation of S397 was required for nuclear cAMP-dependent YAP export from the nucleus and tumor growth suppression. Furthermore, nuclear cAMP signaling inhibited a pro-tumorigenic transcription program that highly correlated with low YAP-dependent gene expression across numerous human melanoma cell lines. Finally, our findings identify nuclear sAC localization as a possible prognostic marker for melanoma and suggest that targeting the nuclear cAMP microdomain may provide a new therapeutic approach for cancer treatment.

## Results

### A nuclear cAMP microdomain is lost upon melanoma

The intraorganellar localization of sAC within mammalian cells is not static and the presence of sAC in the nucleus is associated with early cellular transformation(Desman et al., 2014; Magro et al., 2012). It is well established that sAC is cytoplasmic in benign melanocytes (e.g., nevus) but upon transformation (e.g., melanoma in situ) nearly all melanocytes exhibit pan-nuclear sAC expression(Barnhill et al., 2013; Desman and Barnhill, 2016; Li et al., 2016; Magro et al., 2012; Solky and Zembowicz, 2014). However, whether nuclear sAC expression was maintained during tumor progression remained unknown. Here, we determined whether sAC localization in the nucleus was stable upon tumor progression. Melanoma is an ideal model to investigate changes in protein expression or localization during tumor progression because the association between melanoma depth of invasion and disease progression or prognosis is well characterized(Schadendorf et al., 2018; Shain and Bastian, 2016). We measured sAC localization in a panel of human melanoma biopsy samples (n = 34). sAC was consistently localized in the nucleus of melanoma cells that are present in the epidermis and reticular dermis (e.g., in situ and early invasive disease). In contrast, sAC became absent in the nucleus of melanoma cells as the tumor invaded into the deeper dermis (Figure 1A-E). We measured the tumor depth at which sAC was lost from the nucleus of melanoma cells using Breslow thickness, a standardized method of measuring melanoma progression and assessing risk of metastasis(Abbas et al., 2014; Paek et al., 2007). We found that loss of sAC from the nucleus correlated significantly with melanoma staging (p < 0.001, likelihood-ratio test; Figure 1F). Thus, sAC was not stably localized in the nucleus during melanoma progression and nuclear staining decreased significantly with the depth of tumor invasion (Figure 1G).

**Figure 1.**
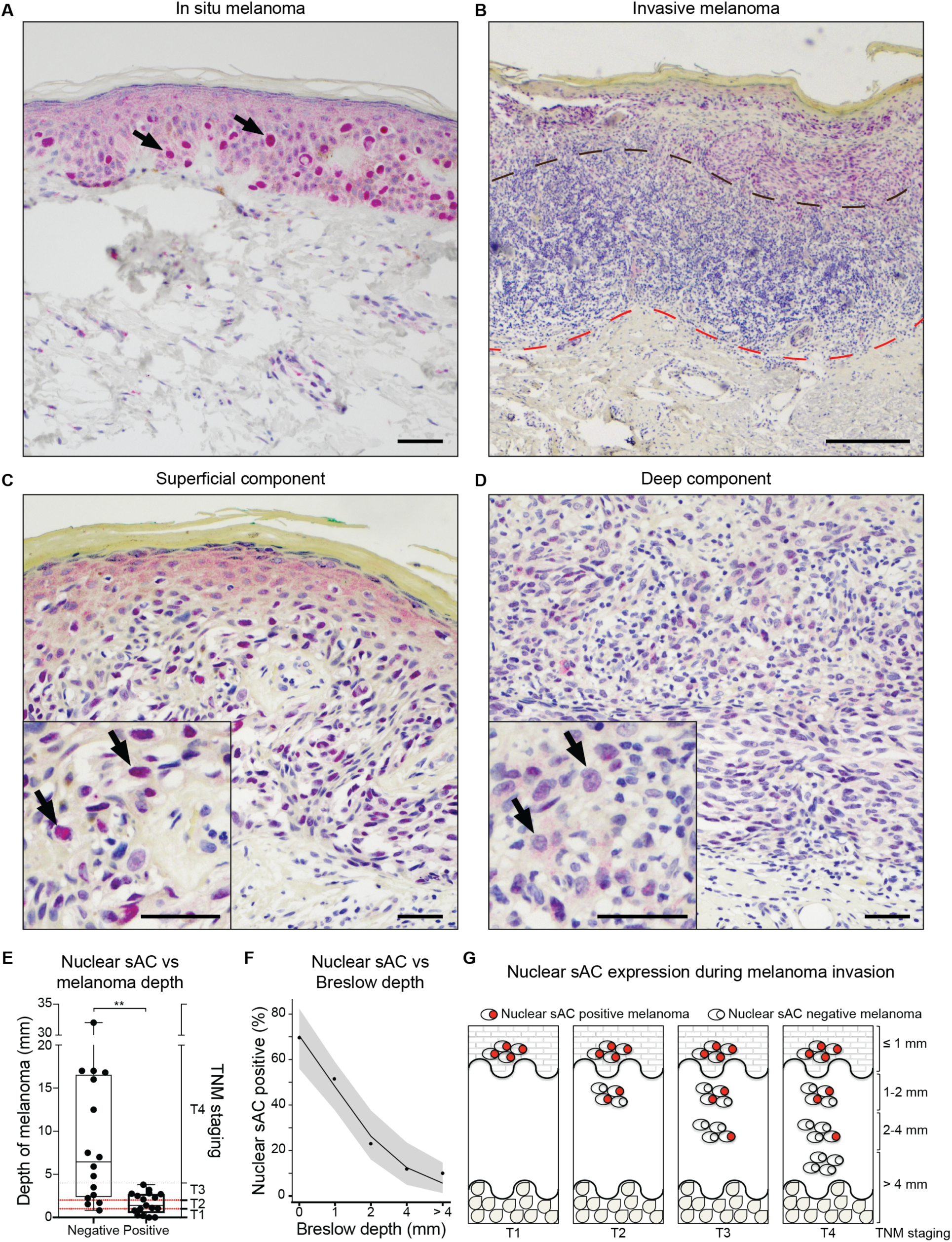
A nuclear cAMP microdomain is lost upon melanoma invasion. A) Subcellular sAC localization in in situ human melanoma; arrows point to examples of nuclear sAC positive cells. Scale bar 50 µm. B) Loss of nuclear sAC upon deep dermal invasion; black line denotes transition from superficial to deep melanoma component, while red line marks the leading edge of the tumor. Scale bar 500 µm. C) Maintenance of nuclear sAC in superficial component of invasive melanoma (indicated by arrows). Scale bars 25 µm. D) Loss of nuclear sAC in the deep dermal component of invasive melanoma (indicated by arrows). Scale bars 50 µm. E) Correlation between melanoma depth and the presence of nuclear sAC positive melanocytes. Welch’s t-test; **, P ≤ 0.01. Box extends from 25^th^ to 75^th^ percentiles, with median indicated inside. Whiskers extend from minimum to maximum value with individual data points shown. F) Increasing Breslow depth correlates with loss of nuclear sAC positive cells in human melanoma. Points represent observed values (%), solid line represents regression model estimates with 95% CI as shaded regions. Significance testing (P < 0.001) was performed using likelihood ratio test. G) Schematic of nuclear sAC expression during different stages of melanoma invasion.

We previously reported that genetic deletion of all sAC regulated microdomains sensitizes fibroblasts and keratinocytes to transformation(Ramos-Espiritu et al., 2016a). In addition, we now show that transplantation of melanocytes with genetic inactivation of sAC (*Adcy10^-/-^*, sAC^KO^) leads to tumor formation in immunodeficient mice while control wild type (WT) immortalized melanocytes fail to grow in mice (Figure S1). Therefore, our data suggest that loss of sAC enhances tumorigenesis in multiple cell types ((Ramos-Espiritu et al., 2016a) and Figure S1). We reasoned that loss of sAC from a specific microdomain increases tumorigenesis.

### Development of a genetic tunable system to investigate cAMP microdomains in vitro and in vivo

To assess the relative contribution of cAMP in different microdomains, we developed a genetic-based, tunable system allowing for the control of cAMP signaling in three spatially distinct microdomains not previously examined in cancer: nucleus (NLS-sAC); cytoplasm (NES- sAC); and mitochondria (mito-sAC) (Figure 2A). These microdomain-targeted constructs were introduced into the sAC^KO^ mouse melanoma cell line (Figure 2B). Doxycycline-induced expression of sAC in each microdomain was rapid (∼1 hour), dose-dependent and led to increased cAMP synthesis (Figure S2A-C). cAMP levels in a cell are tightly regulated and reflect a balance between adenylyl cyclase (AC)-dependent cAMP synthesis and cAMP degradation by PDEs within each microdomain(Cooper and Tabbasum, 2014; Lohse et al., 2017). Consistent with each of these targeted ACs forming PDE-regulated cAMP microdomains, inhibition of PDEs by IBMX led to dramatic increases in cAMP as compared to LacZ controls (Figure S2D). Mitochondrial sAC (mito-sAC) generated cAMP was previously shown to exclusively reside within the mitochondrial matrix, and cytoplasmic cAMP does not penetrate into this microdomain(Acin-Perez et al., 2009a; Acin-Perez et al., 2009b; Valsecchi et al., 2017). However, because cAMP can diffuse across the nuclear pore complex, we wanted to confirm that our NES-sAC and NLS-sAC generated microdomains were distinct. Using an established, genetically-encoded, live-imaging fluorescence resonance energy transfer (FRET)-based sensor of cAMP(Sample et al., 2012b), we confirmed that NLS-sAC-dependent cAMP is localized only to the nucleus and that NES-sAC-dependent cAMP is localized only to the cytoplasm (Figure S2E). Therefore, microdomain-targeted ACs are not overwhelming the normal regulatory control mechanisms of cAMP signaling and thus function biochemically similar to endogenous ACs. Our data demonstrate that we have generated a tunable genetic approach allowing us to test the role of cAMP in multiple intracellular domains. We next asked which microdomain(s) affected tumor cell growth.

**Figure 2.**
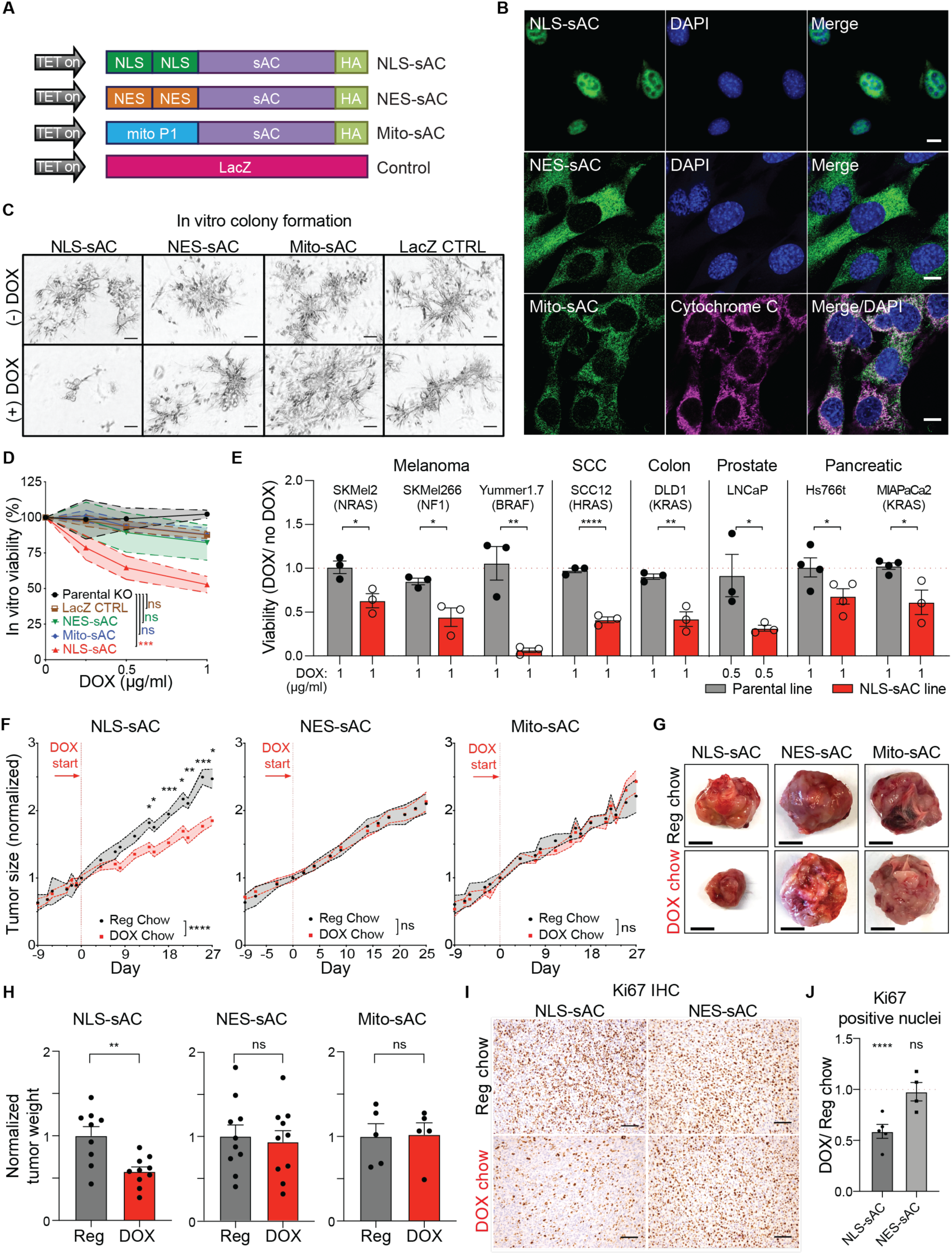
The nuclear cAMP microdomain suppresses tumor growth in vitro and in vivo. A) cDNA schematics of sAC constructs used to study cAMP microdomain biology. NLS, Nuclear Localization Signal; NES, Nuclear Export Signal; mito P1, Mitochondrial P1 Localization Signal; LacZ, Beta galactosidase encoding gene; TET on, doxycycline-activated promoter; HA, hemagglutinin tag. B) Fluorescent light microscopy images of melanoma clones cultured in the presence of doxycycline that show targeted localization of sAC microdomains as detected with anti-HA antibody (green). DAPI (blue) used to label nuclei. Anti-cytochrome c antibody (magenta) used to label mitochondria. Scale bar 10 µm. C) In vitro colony formation by mouse melanoma cells in Matrigel after 7 days +/- doxycycline (DOX; 1µg/ml). Scale bar 100 µm. D) Viability of mouse melanoma colonies formed in vitro in Matrigel, assessed by luminescence-based measurement of ATP content. n=3 or more; error bars, SEM; Two-way ANOVA. E) Viability of human and mouse cancer cell lines following nuclear sAC expression in vitro in Matrigel, assessed by luminescence-based measurement of ATP content. Error bars, SEM; Student’s t-test, n=3. F) In vivo tumor growth of melanoma cells expressing microdomain targeted sAC. Switch to doxycycline (DOX) containing chow is indicated by the arrow and red vertical line. Data normalized to day 0 (day when mice changed to DOX containing chow). Reg, regular chow, gray. DOX, Doxycycline containing chow, red line. Dashed lines, SEM; mixed effect ANOVA, with Sidak correction for multiple comparisons. NLS-sAC, n=15 per cohort; NES-sAC and Mito-sAC n=10 per cohort. G) Examples of gross tumor images from the experiment in (F). Scale bar 1 cm. H) Weight of NLS-sAC, NES-sAC, and Mito-sAC tumors from mice fed regular (Reg, gray bars) or doxycycline containing (DOX, red bars) chow. Weight was normalized within each pair to tumors from the cohort fed regular chow (set to 1). Error bars, SEM; Student’s t-test. I) Microscopic image examples of Ki67 immunohistochemistry (IHC) analysis of tumor sections. Scale bar 100 µm. J) Quantitation of Ki67 positive nuclei in tumor sections from in vivo experiment represented as a fold change over corresponding control cohort. Mean with data points for individual tumors are shown. Error bars, SEM; Student’s t-test. (ns, P > 0.05; *, P ≤ 0.05; **, P ≤ 0.01; ***, P ≤ 0.001; ****, P ≤ 0.0001).

### The nuclear cAMP microdomain suppresses tumor growth in vitro and in vivo

The ability to grow in a Matrigel media is a hallmark of cellular transformation(Benton et al., 2011). Mouse melanoma cell proliferation as three-dimensional colonies in Matrigel was unaffected by LacZ expression, an increase in cAMP signaling in the cytoplasm (NES-sAC) or inside the mitochondria (mito-sAC). In contrast, activation of nuclear cAMP signaling (NLS-sAC) inhibited the growth of melanoma cells in Matrigel (Figure 2C, D). To confirm whether inhibition of tumor cell growth by nuclear cAMP signaling was dependent upon tumor cell type, we established a panel of human and mouse cancer cell lines containing the doxycycline-inducible NLS-sAC cassette (Figure 2A). Similar to our results in sAC^KO^ mouse melanoma cells, activation of nuclear cAMP signaling reduced the growth of multiple human and mouse cancer cell lines in vitro (Figure 2E and Figure S3). Next, we tested whether nuclear cAMP signaling affects tumor progression in mice. NSG mice were implanted with LacZ, NES-sAC, mito-sAC, or NLS-sAC melanoma cells and tumor volume was monitored over time. When tumors reached 1 cm in diameter, mice were randomized to either normal chow or doxycycline containing chow to induce transgene expression. Consistent with our observations in Matrigel, activation of cAMP signaling in the cytoplasm or mitochondria had no effect on tumor progression in mice; similarly, no effect on tumor growth was observed in the LacZ control (Figures 2F-H and Figure S4). However, in contrast, activation of nuclear cAMP signaling led to a significant inhibition of tumor growth as measured by a reduction in tumor diameter, tumor weight and Ki67 nuclear staining (Figures 2F-J and Figure S4B). Our data show that, cAMP signaling in the nucleus, but not in mitochondria or the cytoplasm, suppresses mouse and human tumor cell proliferation suggesting that cAMP has differential effects depending on its localization.

### The nuclear cAMP microdomain uniquely alters chromatin accessibility and inhibits pro-tumorigenic gene expression profiles

The existence of a nuclear cAMP signaling cascade was first reported by our group(Zippin et al., 2003; Zippin et al., 2004a) and has since been confirmed by others(Sample et al., 2012a), but the function of this cAMP microdomain has remained unclear. We posited that differential nuclear cAMP signaling may suppress tumor cell growth by altering gene expression profiles critical for tumor growth. RNA-seq analysis of NES-sAC, mito-sAC, and NLS-sAC expressing melanoma tumors in vivo revealed that activation of cAMP signaling in each microdomain led to significant changes in gene expression (Figure S5A). Whereas nuclear cAMP led to the most potent changes in gene expression among the cAMP microdomains, we were surprised to learn that there was minimal overlap between the genes associated with each microdomain (Figures S5A and S6). These data further confirmed that there is limited cAMP diffusion (Figure S2E and (Acin-Perez et al., 2009b)) between each sAC-defined microdomain and that each of these cAMP microdomains affect distinct gene expression programs. Gene set enrichment analysis (GSEA) of the nuclear cAMP microdomain revealed that nuclear cAMP uniquely led to specific changes in pathways predicted to suppress tumor growth, e.g. downregulation of Myc targets (Figure 3A and S5B). Furthermore, the most significant GSEA pathways upregulated or downregulated by nuclear cAMP (Figure S5B; left and right panel, respectively) compared to the other sAC-defined microdomains confirmed that activation of the nuclear cAMP microdomain led to mRNA changes unique from the other sAC-defined microdomains in tumors in vivo. Thus, our observations show that nuclear, cytoplasmic, and mitochondrial sAC cAMP lead to distinct gene expression profiles in cancer.

**Figure 3.**
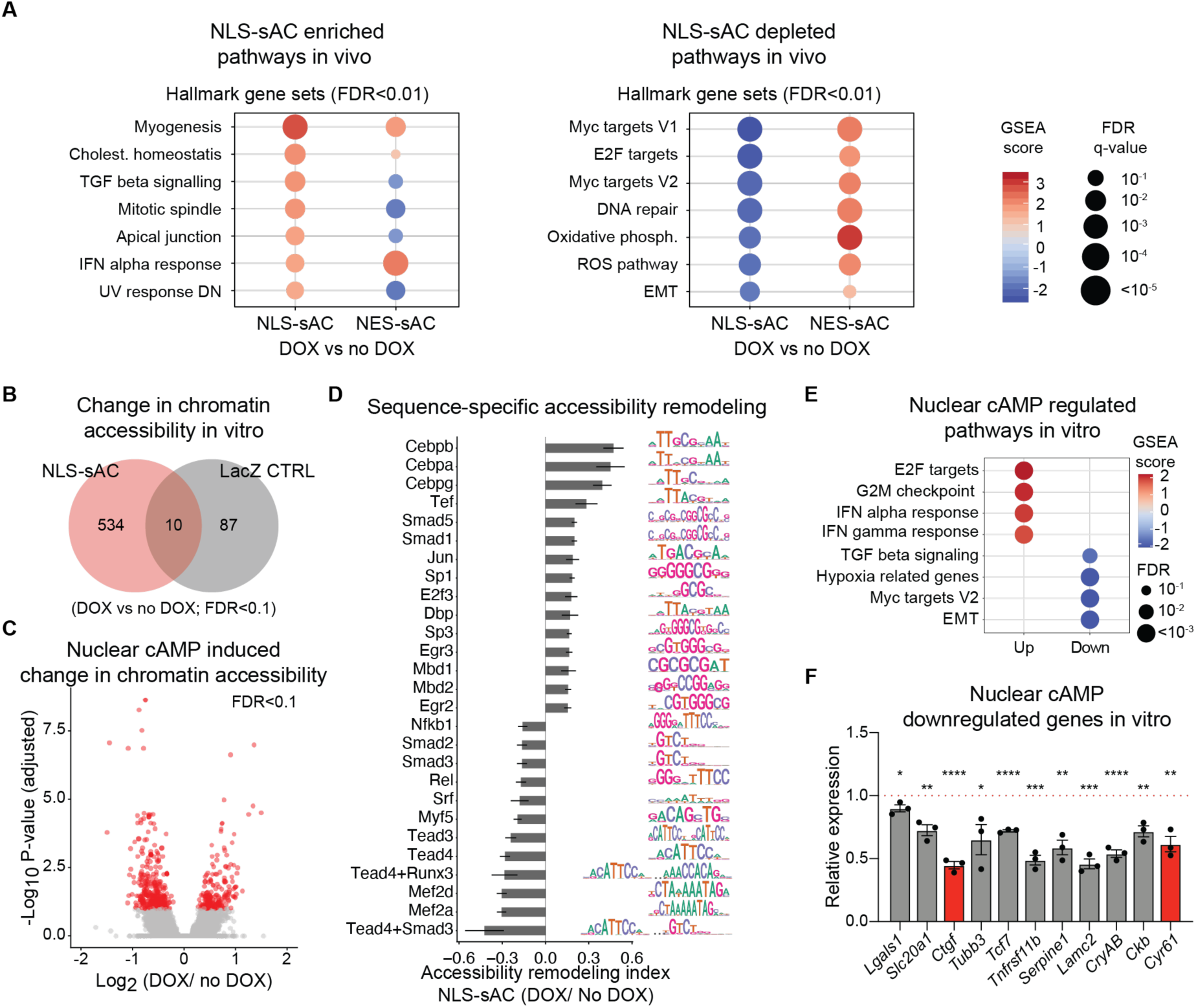
The nuclear cAMP microdomain uniquely alters chromatin accessibility and inhibits pro-tumorigenic gene expression profiles. A) Gene Set Enrichment Analysis of global expression changes in NLS-sAC (DOX) vs control (no DOX) and NES-sAC (DOX) vs control (no DOX) expressing tumors. Upregulated and downregulated pathway panels list gene sets with significant (FDR < 0.01) positive and negative enrichment, respectively, in NLS-sAC vs control. B) Minimal overlap of differential chromatin accessibility following 48h doxycycline treatment (DOX) as compared to no DOX in NLS-sAC and LacZ control cell lines. C) Volcano plot of differential chromatin accessibility of DNA elements in NLS-sAC expressing cells after 48h of doxycycline treatment (DOX) as compared to no DOX. D) Change in transcription factor accessibility remodeling following nuclear cAMP expression. The 27 most significant transcription factors (FDR < 1 x 10^-5^) are reported with their consensus motif sequence logos indicated on the right. Error bars, 95% confidence interval E) Gene Set Enrichment Analysis of the immediate effects of NLS-sAC expression in mouse melanoma cells after 48h of doxycycline treatment. F) RT-PCR confirmation of genes with reduced expression identified by RNA-seq on cells treated with doxycycline for 48h. Hippo pathway genes labeled as red bars. Represented as DOX/no DOX. Mean values are shown. Error bars, SEM. Student’s t-test. (*, P ≤ 0.05; **, P ≤ 0.01; ***, P ≤ 0.001; ****, P ≤ 0.0001).

To better understand the immediate impact of nuclear cAMP on gene expression, we defined the chromatin accessibility regulatory landscape of NLS-sAC expressing melanoma cells by ATAC-seq and RNA-seq in vitro. We constructed a chromatin accessibility atlas across cultured cells consisting of 66,275 ATAC-seq DNA elements and found that activation of nuclear cAMP signaling led to specific and significant changes in chromatin accessibility (Figures 3B, C and S7). We identified 544 DNA elements with differential chromatin accessibility following NLS-sAC expression (FDR < 0.1; DOX vs no DOX), compared to 97 DNA elements in LacZ negative control experiments, suggesting changes in transcription factor (TF) activity upon nuclear cAMP activation. To address this point and considering that chromatin accessibility at TF DNA motifs reflects nucleosome displacement due to TF binding and chromatin remodeler recruitment(Shashikant and Ettensohn, 2019), we developed a computational method to infer the changes in TF accessibility remodeling. We used this approach to study the relative contribution of 330 TFs expressed in these cells to the changes in chromatin accessibility across the 66,275 DNA elements (Figures 3D and S5C). Many of the transcription factors with decreased accessibility remodeling (e.g., TEAD, SMAD, RUNX) are important for tumorigenesis and are regulated downstream of signaling pathways with known roles in cancer(Grannas et al., 2015; Passaniti et al., 2017; Zhou et al., 2016). Among them is the Hippo pathway, an important regulator of cancer cell proliferation, invasion, and apoptosis(Yu et al., 2015b). This pathway was predicted from our ATAC-seq data to be inhibited by nuclear cAMP.

We next performed RNA-seq analysis on tumor cells in vitro (Figure S5D), which confirmed significant correlation between changes in chromatin accessibility of ATAC-seq DNA elements and the expression of genes adjacent to those elements (Figure S5E). GSEA of the changes in gene expression following NLS-sAC induction revealed significant enrichment in pathways that are relevant to tumor biology such as Interferon response, hypoxia, epithelial-to-mesenchymal transition, TGF beta signaling, and Myc signaling (Figure 3E). Many of these pathways are driven by the Hippo-dependent transcription factors identified in our ATAC-seq analysis (Figure 3D). We confirmed expression changes of several cancer-relevant genes identified by RNA-seq, including the reduced expression of the canonical Hippo pathway dependent genes, e.g., *Ctgf* (also known as *Ccn2*) and *Cyr61* (also known as *Ccn1*) (Figure 3F and S5F) by qRT-PCR(Ramos-Espiritu et al., 2016b). Interestingly, RNA-seq analysis from both tumors and cells showed that canonical cAMP-dependent melanocyte gene expression profiles (e.g., *Mitf*) were unaffected by either cytoplasmic, mitochondrial, or nuclear sAC cAMP microdomains. We confirmed that, consistent with previous reports(Goding and Arnheiter, 2019; Hsiao and Fisher, 2014; Johannessen et al., 2013), activation of tmACs at the plasma membrane by GPCR agonists results in increased *Mitf* expression in melanoma cells, whereas induction of nuclear sAC does not (Figure S5G-I). These data taken together show that multiple cAMP microdomains defined by either sAC or tmACs regulate distinct gene expression profiles in melanoma cells and suggest that NLS-sAC may regulate the Hippo signaling pathway.

### The nuclear cAMP microdomain suppresses tumor growth via YAP S397 phosphorylation and export of YAP from the nucleus

The Hippo pathway controls cancer cell proliferation and invasion principally by regulating the activity of the transcriptional coactivators YAP and TAZ (Figure 4A); thus, we reasoned that nuclear cAMP signaling may affect YAP/TAZ activity. Of note, a connection between sAC activity and Hippo signaling has been suggested in recent studies but a mechanistic link between sAC and Hippo signaling has remained unclear(Wang et al., 2020; White et al., 2019). cAMP signaling is reported to regulate YAP/TAZ activity; however, these studies were focused entirely on plasma membrane generated cAMP(Kim et al., 2013). Thus, we next asked if nuclear cAMP signaling represents a previously unrecognized mechanism of YAP and/or TAZ regulation.

**Figure 4.**
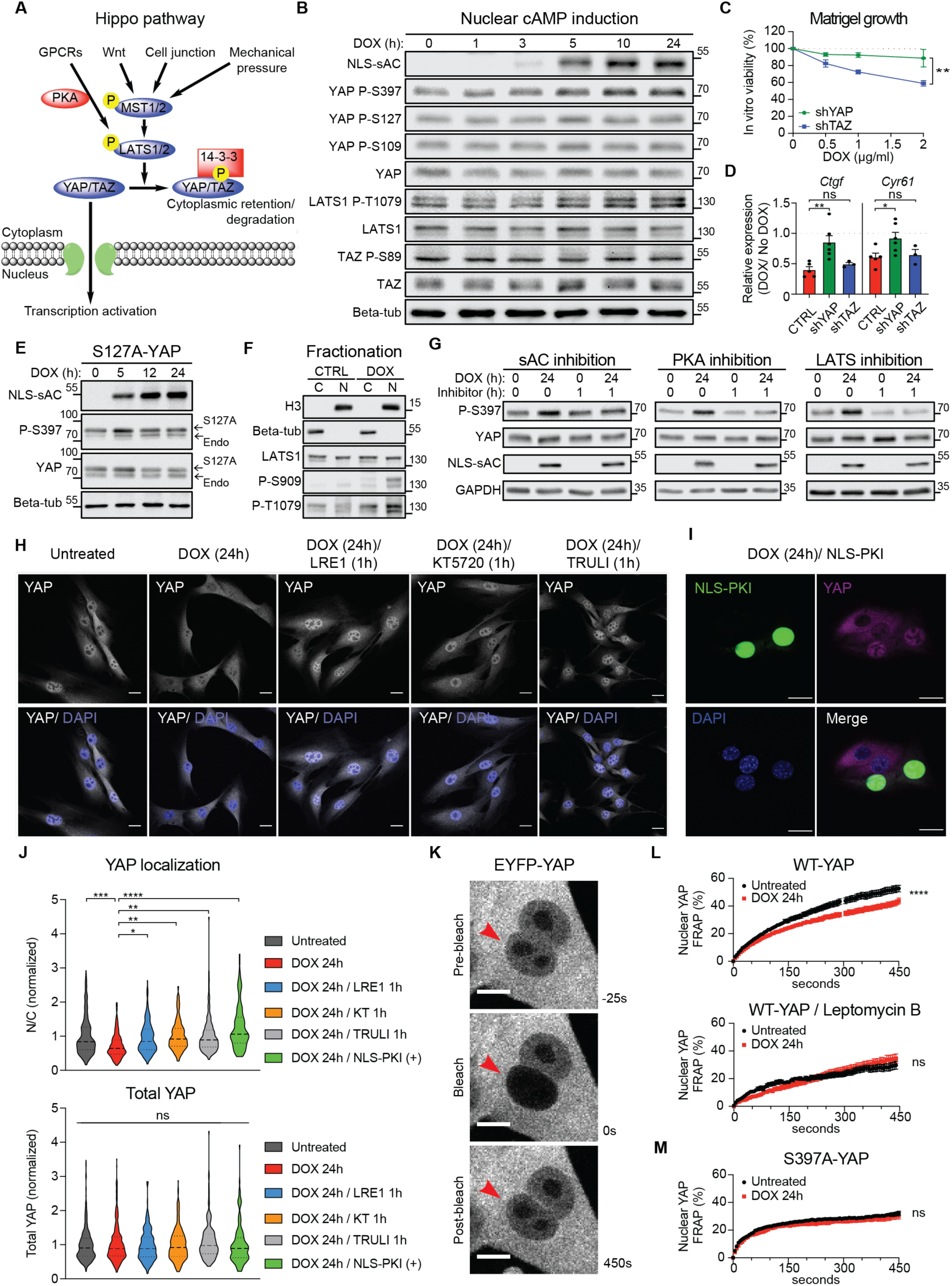
The nuclear cAMP microdomain leads to YAP S397 phosphorylation and YAP nuclear export. A) Schematic of canonical Hippo pathway with key regulators indicated. B) Western blot analysis of nuclear cAMP induced phosphorylation of LATS1, YAP and TAZ shows that only YAP at S397 and LATS1 are phosphorylated. C) Viability of mouse melanoma colonies with either YAP or TAZ knock-down in vitro in Matrigel, assessed by luminescence-based measurement of ATP content. n≥4; error bars, SEM; Two-way ANOVA D) Nuclear cAMP induction by doxycycline leads to reduction of *Ctgf* and *Cyr61* expression (CTRL, red), as measured by qRT-PCR. Knock-down of YAP abolishes nuclear cAMP effect on the expression of these genes (shYAP, green), while TAZ knock-down (shTAZ, blue) has no effect. Expression relative to cells without doxycycline treatment. (n≥4; error bars, SEM; Student’s t-test). E) Phosphorylation of S397 induced by nuclear cAMP does not require S127 phosphorylation (n=3; representative shown). S127A, FLAG-tagged YAP with S127 mutated to Alanine. Endo, endogenous YAP. F) Fractionation of melanoma cells demonstrates the presence of LATS in both the cytoplasmic (C) and nuclear (N) fractions. Active LATS (phosphorylated on both S909 and T1079) is enriched in the nuclear fraction of cells treated with doxycycline (DOX; 24h). Histone H3 and Beta-Tubulin used as markers for nuclear and cytoplasmic fractions, respectively. G) Nuclear cAMP induced phosphorylation of YAP at S397 is abolished by treatment with the sAC inhibitor LRE1, the PKA inhibitor KT5720, and the LATS kinase inhibitor TRULI. H) Microscopic images showing loss of nuclear YAP upon NLS-sAC induction with doxycycline (DOX) and inhibition of YAP nuclear loss by sAC, PKA, and LATS inhibitors (LRE, KT5720, and TRULI, respectively) when added 1 hour before fixing the cells (scale bar 20 µm). I) Microscopic images of YAP (purple) nuclear expression following doxycycline (DOX) treatment of NLS-sAC melanoma cells in the presence or absence of a nucleus-targeted Protein Kinase A inhibitor (NLS-PKI, green). Scale bar 20 µm J) Quantification of YAP localization represented as nuclear to cytoplasmic ratio (N/C, upper panel) following nuclear cAMP induction alone (DOX 24h), or in the presence of sAC inhibitor (LRE1), PKA inhibitors (KT5720 and NLS-PKI), or LATS inhibitor (TRULI); total YAP level (lower panel) remained unchanged under all conditions. Dashed line within each violin plot, Median; doted lines, Quartiles; error bars, SEM; one-way ANOVA with Sidak correction for multiple comparisons. N= 3-6 individual experiments. Total cell number per group: 446 Untreated, 477 DOX 24h, 221 DOX 24h / LRE1 1h, 252 DOX 24h / KT5720 1h, 212 DOX 24h / TRULI 1h, and 143 DOX 24h / NLS-PKI (+). K) Microscopic image examples of FRAP analysis of EYFP tagged YAP. Bleached nucleus is marked by an arrowhead. Scale bar 10 µm. L) FRAP analysis of EYFP-YAP following induction of nuclear cAMP with doxycycline (DOX) in the presence or absence of the nuclear export inhibitor Leptomycin B. (without Leptomycin B: n=3, 47 nuclei total per group; with Leptomycin B: n=2, 13 nuclei total per group; error bars, SEM; mixed effect ANOVA). M) FRAP analysis of EYFP-tagged YAP-S397A showing no changes in nuclear YAP recovery regardless of doxycycline treatment (n=3, 52 untreated nuclei and 32 doxycycline treated nuclei; error bars, SEM; mixed effect ANOVA). (ns, P > 0.05; *, P ≤ 0.05; **, P ≤ 0.01; ***, P ≤ 0.001; ****, P ≤ 0.0001).

The LATS-dependent phosphorylation of specific serine residues is the principal mechanism of regulation for both YAP and TAZ(Zhao et al., 2010). We therefore investigated the effect of nuclear sAC cAMP activation on YAP and TAZ at specific sites of phosphorylation. Activation of nuclear sAC cAMP signaling led to an increase in LATS phosphorylation, and in turn, YAP phosphorylation at serine 397 (YAP-S397) but not at other YAP or TAZ serine residues (Figures 4B and S8A). Activation of cytoplasmic and mitochondrial sAC cAMP microdomains in melanoma cells did not affect YAP or TAZ phosphorylation, suggesting that regulation of YAP-S397 is unique to the nuclear sAC cAMP signaling microdomain (Figure S8B-D). Consistent with published reports, GPCR-regulated cAMP in these cells led to the phosphorylation of YAP at multiple serine residues, as well as increased phosphorylation of TAZ at S89 (Figure S8E-F)(Yu et al., 2012). Thus, plasma membrane and nuclear cAMP microdomains, albeit differently, both regulate Hippo signaling by altering YAP phosphorylation, whereas cytoplasmic and mitochondrial sAC cAMP microdomains have no effect on YAP. We showed earlier (Figures 2E and S3) that nuclear cAMP inhibits the growth of several cancer cell lines; thus, we looked at their YAP-S397 phosphorylation status and found that S397 phosphorylation is indeed induced by nuclear cAMP signaling across multiple cancer cell types (Figure S8G).

Even though we do not observe any changes in TAZ phosphorylation following nuclear cAMP activation (Figures 4B and S8A), recent studies have suggested that sAC may affect TAZ activity, perhaps by changing TAZ expression(Wang et al., 2020). To firmly establish the necessity of YAP for nuclear sAC tumor suppression, we employed shRNAs to knock down YAP or TAZ in melanoma cells (Figure S9A). Knockdown of TAZ had no effect on nuclear sAC inhibition of tumor cell growth in Matrigel (Figure 4C) or Hippo-dependent gene expression (Figure 4D). In contrast, knock down of YAP prevented nuclear sAC inhibition of tumor cell growth in Matrigel (Figure 4C) and Hippo-dependent gene expression (Figure 4D).

Previous studies report that YAP is phosphorylated at serine residues in a specific order with S127 occurring before S397 and that phosphorylation at S127 may be required for S397 phosphorylation(Zhao et al., 2010). However, these studies were focused on cytoplasmic regulation of YAP. To determine whether S127 phosphorylation is required for nuclear cAMP regulation of S397 phosphorylation, we overexpressed S127A-YAP in melanoma cells. Nuclear cAMP signaling induced S397 phosphorylation of S127A-YAP protein; thus, S127 phosphorylation is not required for S397 phosphorylation in the nuclear cAMP microdomain (Figures 4E and S9B).

We performed cellular fractionation experiments on NLS-sAC melanoma cells and, consistent with published reports(Li et al., 2014), we found LATS kinase localized within the nucleus of NLS-sAC expressing melanoma cells (Figure 4F). Furthermore, activation of nuclear cAMP signaling induced the phosphorylation of LATS kinase within the nucleus (Figure 4F). Thus, the nuclear cAMP signaling cascade leads to both nuclear LATS activation and nuclear YAP S397 phosphorylation. There are two LATS kinases expressed in mammalian cells, LATS1 and LATS2. To test if nuclear cAMP-evoked signaling shows any bias towards LATS1 or LATS2, we used shRNA to knock-down the LATS1 or LATS2 in NLS-sAC melanoma cells (Figure S9C). The cells were then treated with doxycycline to induce nuclear cAMP, and we observed that the knock-down of a single LATS kinase significantly reduced S397 phosphorylation, but did not abolish it completely (Figure S9D). This suggests that the nuclear cAMP domain may act through LATS1 and LATS2.

NLS-sAC expression could induce LATS and YAP phosphorylation either through protein-protein interactions or via the generation of cAMP and the activation of cAMP-effector proteins. YAP-S397 phosphorylation by nuclear sAC was abolished by pharmacologic inhibition of sAC thus cAMP generation within the nucleus is required (Figures 4G and S9E). LATS can be phosphorylated by the cAMP-effector protein PKA(Dasgupta and McCollum, 2019; Yu et al., 2013), and PKA holoenzyme is also present within the nucleus(Zippin et al., 2004a); therefore, we tested whether PKA activity was required for NLS-sAC-dependent activation of nuclear LATS. Pharmacologic inhibition of PKA blocked NLS-sAC induced YAP-S397 phosphorylation thus PKA activity is required (Figures 4G and S9E). We also confirmed that LATS phosphorylation was blocked by inhibitors against sAC and PKA (Figure S9F). To confirm that the NLS-sAC-induced YAP S397 phosphorylation requires active LATS kinase, we incubated cells with the LATS1/2 specific inhibitor TRULI(Kastan et al., 2021). Incubation with TRULI led to complete suppression of NLS-sAC-induced YAP S397 phosphorylation (Figures 4G and S9E). These data confirm that NLS-sAC stimulates a nuclear cAMP microdomain leading to both PKA and LATS activation and YAP S397 phosphorylation.

Phosphorylation of YAP can lead to its export from the nucleus and/or its degradation(Dasgupta and McCollum, 2019; Zhao et al., 2010; Zhao et al., 2007). We did not observe any change in YAP protein level following nuclear cAMP activation (Figures 4B and S8A). We next compared the effect of NLS-sAC activation and GPCR activation on YAP protein level in the presence of the protein translation inhibitor cycloheximide. Whereas NLS- sAC activation did not reduce YAP protein levels relative to control cells, GPCR induced cAMP synthesis did lead to a reduction in total YAP levels relative to control cells (Figure S9G). Thus, NLS-sAC activation does not affect YAP stability. Therefore, we next asked if phosphorylation of nuclear YAP leads to a change in YAP localization within the nucleus. Indeed, activation of nuclear cAMP signaling led to decreased detection of YAP in the nucleus and was associated with an increase in cytoplasmic YAP (Figure 4H). This change in YAP subcellular distribution was blocked by pharmacologic inhibitors of sAC, PKA, and LATS (Figure 4H). Pharmacologic inhibitors of PKA can block both cytoplasmic and nuclear PKA activity. To confirm that nuclear PKA activity was required for the change in subcellular YAP localization by nuclear cAMP, we generated NLS-sAC melanoma cells that express NLS-PKI-GFP(Billiard et al., 2001). We then compared a mixed population of NLS-PKI-GFP positive and negative NLS-sAC cells and found that NLS-PKI-GFP expression blocked nuclear cAMP-induced reduction of YAP in the nucleus (Figure 4I). The quantitation of nuclear and cytoplasmic YAP expression, and total YAP detection across multiple conditions is summarized in Figure 4J. Reduction of nuclear YAP levels following nuclear cAMP activation suggests that the nuclear cAMP cascade either prevents YAP import and/or induces YAP export. To address this question, we generated NLS- sAC melanoma cells expressing EYFP-YAP(Ege et al., 2018) and performed fluorescence recovery after photobleaching (FRAP) analysis (Figure 4K). Nuclear sAC expression did not delay the immediate recovery after photobleaching but did prevent the recovery of YAP to wild type levels (Figure 4L). These data suggested that the initial import of YAP into the nucleus was not prevented by nuclear sAC. We next repeated these experiments in the presence of leptomycin B, an inhibitor of nuclear export. Inhibition of nuclear export blocked the reduction of nuclear YAP levels following nuclear cAMP activation suggesting that nuclear cAMP signaling promotes YAP export from the nucleus (Figure 4L). Finally, we asked whether YAP S397 phosphorylation was required for NLS-sAC-induced export of YAP from the nucleus. We performed site-directed-mutagenesis on the EYFP-YAP cDNA to generate a EYFP-YAP-S397A mutant. Overexpression of EYFP-YAP-S397A prevented NLS-sAC-induced export of YAP from the nucleus (Figure 4M). Thus, our data suggests that nuclear sAC activation does not affect YAP protein stability and leads to the export of YAP from the nucleus via phosphorylation of S397. Since nuclear cAMP-induced export of YAP requires S397 phosphorylation, we next examined whether YAP-S397 phosphorylation was necessary for nuclear cAMP-dependent suppression of tumor proliferation.

Using our NLS-sAC melanoma cell lines, we established sub-clones expressing either wild type YAP (WT-YAP), YAP-S397A (S397A), YAP-S127A (S127A), or YAP-5SA (5SA; non-phosphorylatable YAP mutant with five key serine residues mutated to alanine, including S397 and S127) (Figures 4E and S10A)(Zhao et al., 2010; Zhao et al., 2007). Similar to parental NLS-sAC cell lines, nuclear cAMP signaling inhibited the proliferation of both WT-YAP and S127A overexpressing melanoma cells in vitro (Figure 5A). In contrast, S397A and 5SA melanoma cells were insensitive to nuclear cAMP signaling, whether assessed by suppression of proliferation in Matrigel or mice (Figures 5A, B and S10B, C). Of note, S397A and 5SA melanoma cells grew at a faster rate as compared to WT-YAP melanoma cells in vivo. The NLS- sAC induced reduction in the Ki67 proliferation marker (Figure 2I, J) was abolished by expression of S397A and 5SA YAP in these cells (Figure S10D, E). Thus, YAP-S397 phosphorylation is necessary for nuclear cAMP-dependent suppression of tumor proliferation.

**Figure 5.**
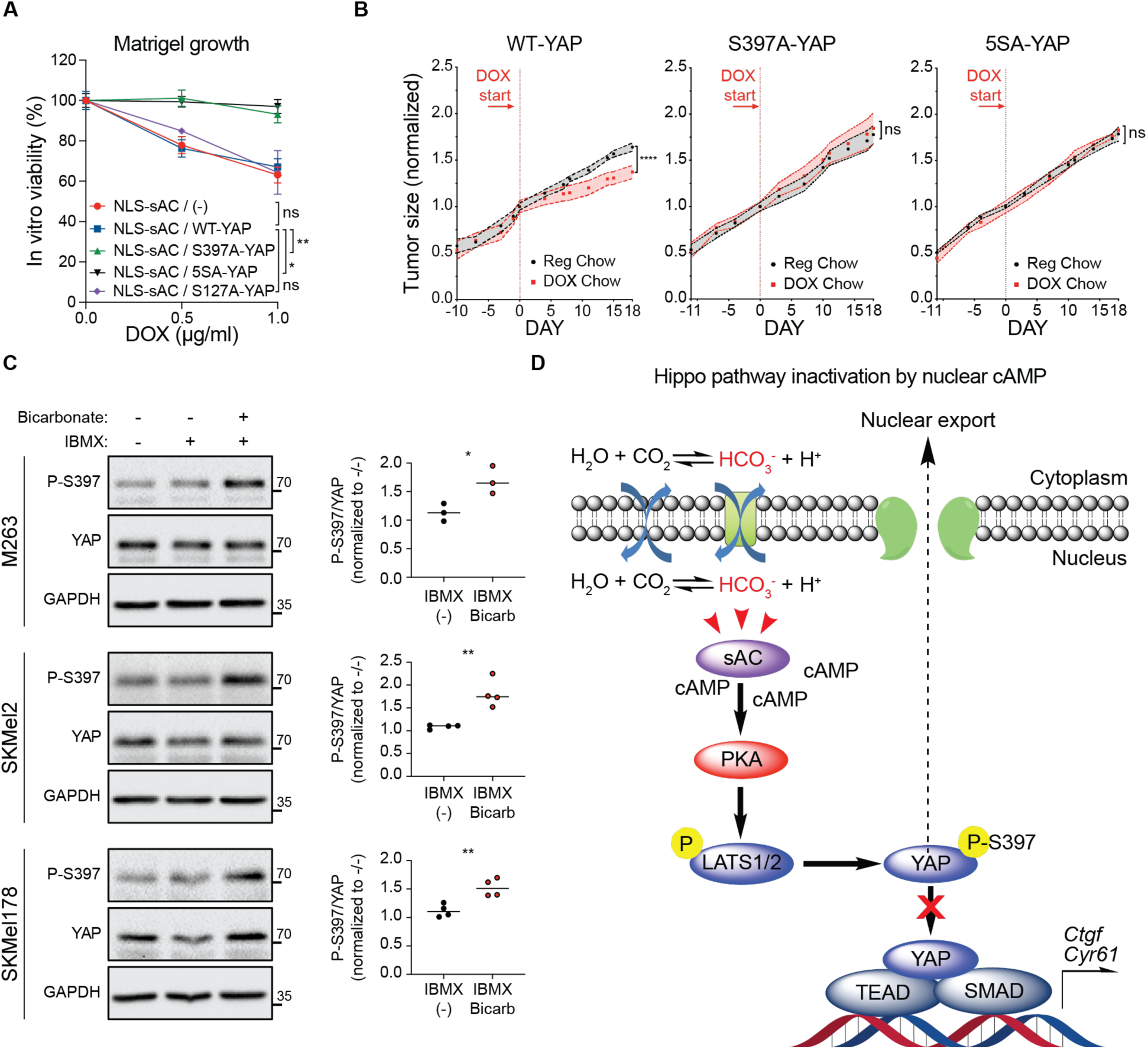
Bicarbonate responsive nuclear sAC inhibits tumor growth through Hippo pathway inactivation. A) In vitro growth of colonies in Matrigel assessed by luminescence-based measurement of ATP content of the parental (-) NLS-sAC melanoma line or melanoma cells overexpressing either wild type (WT)-YAP or mutant YAPs (S127A, S397A or 5SA). n=6; error bars, SEM; ANOVA. B) Loss of in vivo tumor growth inhibition by nuclear cAMP in melanoma cells that express either S397A- or 5SA-YAP mutants. Switch to doxycycline containing chow (DOX chow, red) is indicated by the arrow and red line. Reg, regular chow, gray; Error bars, SEM; mixed effect ANOVA, with Sidak correction for multiple comparisons; n=5 per cohort, except 5SA- YAP/DOX chow n=4. C) Left panel, representative Western blots showing YAP S397 phosphorylation in human melanoma lines incubated in control media or in the presence of 50 µM IBMX with or without bicarbonate ion (HCO_3_^-^), an agonist of endogenous sAC, for 30 mins. Right panel, quantification of P-397 band volumes normalized to total YAP band intensity expressed as the fold change of bicarbonate+IBMX (+,+) relative to IBMX alone (-,+). n≥3. Student’s t-test. D) Model of Hippo pathway inhibition by nuclear cAMP. (ns, P > 0.05; *, P ≤ 0.05; **, P ≤ 0.01; ***, P ≤ 0.001; ****, P ≤ 0.0001).

Tumors grow in a relatively acidic microenvironment and are known to be metabolically active(Liberti and Locasale, 2016; Marino et al., 2012). This unique microenvironment is known to promote cancer growth and invasion(Liberti and Locasale, 2016; Marino et al., 2012). YAP is known to respond to products of metabolism and Hippo signaling can affect multiple metabolic pathways(Koo and Guan, 2018; Zhang et al., 2018). sAC is a pH/metabolic sensor capable of responding to changes in the CO_2_/bicarbonate/pH level in cells, which is in constant equilibrium(Chang and Oude-Elferink, 2014; Zippin et al., 2013). Thus, we reasoned that sAC may represent an additional link between pH and/or metabolism and Hippo signaling. To determine whether the nuclear cAMP microdomain was responsive to changes in intracellular bicarbonate levels, we grew human melanoma cells in pH 7.2 stabilized media deficient in bicarbonate and under ambient CO_2_ conditions, and then stimulated cells with bicarbonate. Bicarbonate stimulation led to activation of endogenously-encoded sAC, and immediate phosphorylation of YAP at S397 (Figure 5C) in multiple human melanoma cell lines. Thus, the local concentration of bicarbonate can regulate Hippo signaling via nuclear sAC (Figure 5D).

### Nuclear cAMP- and YAP-dependent signaling are inversely correlated in human melanoma

YAP signaling leads to a defined gene expression pattern in human melanoma(Zhang et al., 2020b). We compared changes in RNA expression following activation of YAP signaling by overexpression of YAP-5SA in the human melanoma cell line MeWo(Zhang et al., 2020b) to the changes in RNA expression induced by NLS-sAC in our mouse melanoma cells grown both in vitro and in vivo. NLS-sAC-dependent gene expression in mouse melanoma cells grown in vitro (P = 1.08e-12) or in mice (P < 2e-16) was inversely correlated with YAP-dependent gene expression in MeWo cells (Figure 6A). To validate this finding, we expanded our analysis to other human melanoma cell lines. We created three distinct gene sets for NLS-sAC and YAP, each enriched in genes that are strong indicators of NLS-sAC and YAP activities. We used these genes with two complementary single-sample gene set enrichment approaches to infer NLS-sAC and YAP activities across 45 melanoma cell lines from the Cancer Cell Line Encyclopedia (CCLE) (see Methods). Regardless of the gene set or enrichment approach used, NLS-sAC and YAP activity signatures were anti-correlated across the melanoma cell lines (Figure 6B).

**Figure 6.**
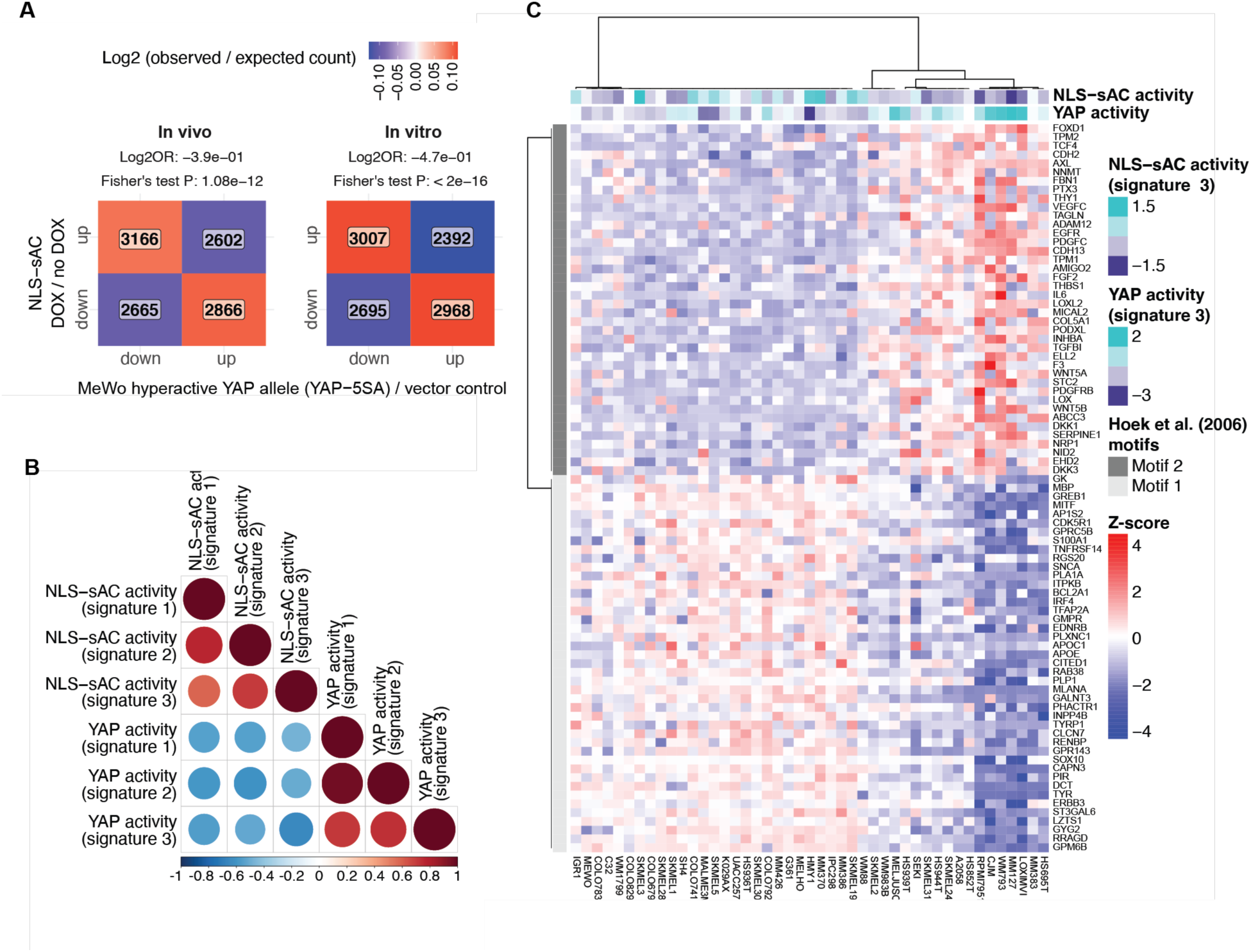
Nuclear cAMP- and YAP-dependent signaling are inversely correlated in human melanoma. A) Heatmaps comparing the relative up-regulation and down-regulation of genes in hyperactive YAP expressing MeWo cells (rows) versus NLS-sAC (columns) expressing in vivo mouse melanoma tumors (left panel) and in vitro mouse melanoma cell lines (right panel). The number of genes observed in each quadrant is indicated. B) Spearman correlation between inferred NLS-sAC and YAP activities across 45 melanoma cell lines from the Cancer Cell Line Encyclopedia (CCLE). Three approaches were used to derive the activity signatures, sigs. 1-3 (see methods). C) Heatmap showing z-score transformed mRNA expression values for melanocytic and neural crest differentiation signature genes (motif 1) and TGFβ-like signal signature genes (motif 2), defined by Hoek et al. (2006), in 45 melanoma cell lines (columns). Inferred NLS-sAC and YAP activities in each cell line are indicated at the top.

Hoek et al. previously identified two groups of genes whose RNA expression distinguished melanoma cells based on their differentiation state and invasion potential(Hoek et al., 2006). In melanoma cell lines from the CCLE, YAP activity was shown to be associated with the differential expression of the marker genes identified by Hoek et al.(Zhang et al., 2020b). To examine how NLS-sAC activity relates to YAP, we used consensus clustering to segregate melanoma cell lines from the CCLE into two groups based on the RNA expression of genes defined by Hoek et al. One group was characterized by higher NLS-sAC activity and lower YAP activity (motif 1). Conversely, the other group exhibited lower NLS-sAC activity and higher YAP activity (motif 2) (Figure 6C). Altogether, these data suggest that NLS-sAC activation inhibits YAP signaling in human melanoma.

## Discussion

For many years, it was unclear how a single second messenger could lead to such disparate effects in cancer. By taking a microdomain approach, we have uncovered a previously unappreciated nuclear cAMP microdomain that functions as a tumor suppressor. The Hippo pathway is critical for numerous functions in cancer cells(Zanconato et al., 2019) and while cAMP is known to affect Hippo signaling, the specific sources of cAMP have remained largely undefined. Previous work has almost exclusively focused on understanding the regulation of YAP and TAZ in the cytoplasm; however, the main function of YAP and TAZ as transcriptional co-activators occurs in the nucleus. Interestingly, proteins that regulate YAP and TAZ, such as LATS, are known to be present in the nucleus and nuclear LATS activation is reported(Li et al., 2014). We now reveal a previously unappreciated role of nuclear cAMP as an inhibitor of nuclear YAP (Figure 5D). Regulation of YAP by the nuclear cAMP microdomain is distinct from GPCR activated plasma membrane cAMP microdomains, which induce YAP phosphorylation on multiple serine residues (Figure S8E, F) and lead to YAP degradation (Figure S9G)(Kim et al., 2013; Yu et al., 2013). In contrast, nuclear cAMP signaling leads to nuclear LATS-activation and phosphorylation of YAP solely at S397. Phosphorylation of YAP at S397 in the nucleus does not affect YAP protein stability but instead leads to nuclear export (Figures 4B, H-M and S8A). Finally, our data suggest that activation of nuclear cAMP signaling inhibits YAP-dependent signaling in human melanoma (Figure 6). The mechanisms that facilitate the nuclear cAMP-specific phosphorylation of YAP at S397 and its export from the nucleus remain important questions.

During this investigation, we have established that sAC defines at least three distinct cAMP microdomains capable of altering gene expression. Furthermore, we identify at least two distinct cAMP signaling cascades that regulate Hippo signaling: a GPCR-regulated tmAC cascade and a bicarbonate-regulated, nuclear sAC cascade. Thus, our data supports the existence of multiple spatially distinct, differentially regulated cAMP microdomains defined by unique sources of cAMP leading to disparate effects in tumor cells. We anticipate that the investigation of individual cAMP microdomains will identify additional microdomain specific cAMP signaling cascades in both tumor and benign cells.

The existence of multiple spatially distinct sAC microdomains in tumor cells may explain why global genetic deletion or mutation of sAC is not favored during tumorigenesis. Whereas nuclear sAC regulates tumor cell growth via Hippo signaling, mitochondrial sAC controls the electron transport chain and cytoplasmic sAC is suggested to control apoptosis(Acin-Perez et al., 2009a; Ladilov and Appukuttan, 2014; Wang et al., 2020). Thus, distinct sAC cAMP microdomains may have differential effects in cancer. Whereas sAC is encoded by a single gene *ADCY10* and most screens for oncogenes and tumor suppressors focus on sequencing approaches, it may not be surprising that alterations in *ADCY10* have not risen to the level of significance. Our data suggest that a comprehensive imaging (e.g., IHC) study across multiple tumor types at different stages may reveal new mechanisms of cAMP signaling in cancer.

It is well-established that cAMP signaling leads to specific changes in gene expression. However, we now reveal that multiple distinct cAMP-mediated pathways can lead to non-overlapping gene expression changes in the same cell. This is significant because it is common practice to associate specific gene expression profiles with cAMP signaling. For example, in melanocytes, the expression of the lineage-defining transcription factor MITF and genes induced by MITF are used as markers for cAMP-dependent gene expression(Goding and Arnheiter, 2019; Johannessen et al., 2013). We now reveal that MITF is only regulated by a subset of cAMP microdomains in melanocytes. This is consistent with our previous report where sAC inhibition in human and mouse melanocytes did not affect MITF protein level(Zhou et al., 2018). Thus, MITF-dependent gene expression is not reflective of all cAMP signaling pathways in melanocytes. Furthermore, our data demonstrate that certain reagents used to study cAMP, which simultaneously activate or inhibit multiple cAMP microdomains, may lead to confounding results. This new fundamental appreciation of the complexities of cAMP-dependent gene expression are likely relevant for numerous cancer cell types and tissues.

It remains unknown what controls nuclear localization of sAC in human cancers and it will be important to establish which upstream signals affect the translocation of sAC into and out of the nucleus. We provide evidence for possible upstream signals important for the regulation of sAC when present in the nucleus. sAC is regulated by changes in cellular pH and metabolism(Chen et al., 2000; Zippin et al., 2013; Zippin et al., 2001), which are known to influence tumor proliferation and metastasis(Spencer and Stanton, 2019; Zhu and Thompson, 2019). Intra- and extracellular pH and metabolism affect intracellular bicarbonate levels(Lee and Hong, 2020) and sAC activity is known to reflect changes in intracellular bicarbonate levels. We now show that changes in intracellular bicarbonate levels via sAC activity lead to YAP phosphorylation at S397 (Figure 5C). Whether additional metabolic- and/or pH-dependent signals regulate Hippo signaling via sAC is an important question. Previous reports revealed that YAP signaling is affected by extracellular pH via the sensing of protons by GPCRs(Tao et al., 2016; Zhu et al., 2015; Zhu et al., 2016). It will be important to compare how changes in extracellular pH via GPCRs and intracellular pH via sAC affect Hippo signaling.

Whereas nuclear sAC localization is associated with early cellular transformation(Desman et al., 2014; Magro et al., 2012), nuclear sAC localization is not static. Multiple studies have shown that nuclear sAC staining is reflective of early transformation of melanoma in humans and anti-sAC antibodies are used as a diagnostic marker(Barnhill et al., 2013; Desman and Barnhill, 2016; Li et al., 2016; Magro et al., 2012; Solky and Zembowicz, 2014). Similar to other established tumor suppressors in melanoma, such as p16(Adelman et al., 2020; Lade-Keller et al., 2014; Zeng et al., 2018), the tumor suppression mechanism by nuclear cAMP is lost when tumor cells invade. Our data suggests that nuclear sAC staining may also be reflective of tumor invasion risk and/or metastasis. Thus, sAC staining may provide both diagnostic(Barnhill et al., 2013; Desman and Barnhill, 2016; Desman et al., 2014; Magro et al., 2012; Rosko et al., 2017) and prognostic information for cancer. Further investigation is required to determine whether sAC immunolocalization may serve as a prognostic marker.

Hippo signaling plays a critical role in the proliferation, differentiation, and apoptosis of nearly every cell and tissue type(Dasgupta and McCollum, 2019; Yu et al., 2015b). Previous work has almost exclusively focused on the regulation of the Hippo signaling proteins YAP and TAZ in the cytoplasm. We have identified a previously unappreciated regulatory pathway of Hippo signaling that is distinct from established mechanisms. Nuclear cAMP-dependent regulation of Hippo signaling occurs in both mouse and human cells derived from a wide range of tissue types. Furthermore, we find that activation of nuclear cAMP signaling is strongly associated with the inactivation of YAP signaling in human melanoma (Figure 6). Therefore, nuclear cAMP-dependent regulation of Hippo signaling is likely to have broad-reaching and fundamental effects on a variety of cells, tissue, and organs.

Multiple members of the cAMP pathway are druggable, and in many cases isoform specific activators and inhibitors for adenylyl cyclases, cAMP effector proteins, and PDEs are established and/or FDA approved(Baillie et al., 2019; Dessauer et al., 2017; Maurice et al., 2014; Ramos-Espiritu et al., 2016b; Wiggins et al., 2018). However, pharmacologic targeting of specific cAMP microdomains remains an untapped therapeutic approach for cancer. We predict that pharmacologic activation of the nuclear cAMP microdomain by either activating the nuclear specific adenylyl cyclase or by inhibiting the nuclear specific PDEs may provide a new therapeutic approach for the treatment of cancer.

## Author Contributions

M.M.D, A.S.D, G.D., I.R.W., T.M., E.P., and J.H.Z. designed the experiments. M.M.D, A.S.D., G.D., R.B., M.R., J.B., M.Y., J.Y., and R.A. generated the Figures. J.Z.W., A.V., K.A., and E.K. generated critical reagents or assisted with in vivo tumor experiments. M.M.D., O.E., T.M., E.P., I.R.W., and J.H.Z. wrote the manuscript with all authors providing feedback.

## Data and materials availability

All unique/stable reagents generated in this study are available from the Lead Contact with a completed Materials Transfer Agreement. Data pertaining to sequencing studies used in this work will be made publicly available prior to publication.

## Methods

### Clinical data

Post-diagnostic archival formalin-fixed paraffin-embedded tissue blocks and H&E slides from a total of 34 primary cutaneous malignant melanomas were retrieved from the IRB-approved melanoma database (IRB# 16-00816) within the Department of Pathology at the Mount Sinai Hospital. Case annotation of patient age, sex, tumor location, and AJCC pTNM stage were provided before histopathological examination (see table below). Cases were evaluated by a board certified dermatopathologist (GTD) for diagnostic confirmation of malignant melanoma, tumor subtype, Breslow thickness, Clark level, ulceration, and radial and vertical growth phases.

Five micrometer thick unstained tissue sections containing tumor (N = 34) were created and stained under IRB approval (IRB# 16-00816) for R21 (anti-sAC, CEP Inc., 1:1000) using an automated Leica-Bond stainer platform. The percentage of tumor cells with nuclear labeling within the epidermis and dermis were recorded. R21 nuclear expression was then evaluated at each Clark and AJCC-relevant Breslow thickness. Regions with 10% or greater nuclear staining were considered positive and regions with less than 10% nuclear staining were considered negative. Metastatic tumors were stained when available (N=10). Metastasis data for all cases extends for at least 5 years since initial skin biopsy was performed.

| Primary cutaneous malignant melanoma cases |  |  |  |  |  |
| --- | --- | --- | --- | --- | --- |
| Case | Patient Age | Sex | Location | Subtype | AJCC pTNM 8<sup>th</sup> Edition |
| 1 | 62 | M | Toe | Nodular | pT4b, N2a, M0 |
| 2 | 60 | M | Flank | Nodular | pT4b, N3c, M0 |
| 3 | 81 | F | Toe | AL | pT3a, N1a, M0 |
| 4 | 23 | F | Shoulder | Nodular | pT4b, N0, M0 |
| 5 | 71 | M | Back | Nodular | pT4b, N1c, M0 |
| 6 | 77 | M | Chest | Nodular | pT4b, N2c, M0 |
| 7 | 84 | M | Thigh | SS | pT3b, N0, M0 |
| 8 | 43 | M | Arm | SS | pT2a, N0, M0 |
| 9 | 86 | F | Back | Nodular | pT4a, N0, M0 |
| 10 | 86 | M | Scalp | SS | pT3b, N0, M0 |
| 11 | 73 | M | Back | Nodular | pT3b, N1a, M0 |
| 12 | 53 | M | Back | SS | pT1b, N0, M0 |
| 13 | 71 | M | Face | LM | pTis, Nx, Mx |
| 14 | 70 | F | Face | LM | pTis, Nx, Mx |
| 15 | 86 | M | Ear | LM | pT3b, N0, M0 |
| 16 | 27 | F | Calf | SS | pT2a, N0, M0 |
| 17 | 50 | F | Back | SS | pT2a, N0, M0 |
| 18 | 87 | M | Ear | LM | pTis, Nx, Mx |
| 19 | 70 | M | Toe | AL | pT4b, N0, M0 |
| 20 | 62 | F | Foot | AL | pT3a, N0, M0 |
| 21 | 90 | F | Thigh | Nodular | pT4b, N3c, M0 |
| 22 | 86 | F | Forehead | LM | pT1b, N0, M0 |
| 23 | 58 | M | Cheek | LM | pT1a, Nx, Mx |
| 24 | 70 | M | Scalp | LM | pT3a, N0, M0 |
| 25 | 67 | M | Scalp | Nodular | pT4b, N3c, M0 |
| 26 | 28 | F | Arm | Nodular | pT2a, N0, M0 |
| 27 | 86 | F | Heel | AL | pT1a, N0, M0 |
| 28 | 47 | M | Chest | SS | pT3a, N0, M0 |
| 29 | 70 | F | Arm | Nodular | pT2a, N0, M0 |
| 30 | 86 | M | Foot | AL | pT3b, Nx, M0 |
| 31 | 67 | M | Arm | Nodular | pT3b, N0, M1a |
| 32 | 31 | F | Cheek | LM | pT1a, Nx, Mx |
| 33 | 53 | F | Foot | AL | pT4b, N3c, M0 |
| 34 | 75 | F | Arm | Nodular | pT3b, N0, M0 |
| F, female; M, male; AL, Acral Lentiginous; LM, Lentigo Maligna; SS, Superficial Spreading |  |  |  |  |  |

### Animal studies

All procedures involving mice were performed with approval from the Weill Cornell Medical College Institutional Animal Care and Use Committee and were performed in compliance with institutional guidelines. Mice were maintained in ventilated cages, on a standard rodent diet of chow or doxycycline diet (Global 2018 base with 625 mg/kg doxycycline hyclate; Teklad, #TD.01306) as indicated, and water ad libitum. Experiments were performed starting at 6-10 weeks of age on a sex-mixed cohort of in-house bred mice. Tumor formation by wild type (sAC FF) and *Adcy10^-/-^* (sAC KO) melanocytes (n=10 per cohort) was performed by injecting into both flanks 5,000,000 cells in 50 µL of 25% Matrigel Basement Membrane Matrix (Corning, #354248). Mice were observed every 3-4 days for tumor formation and tumor size was measured. Mice were euthanized when tumors reached 2cm. For studies of the effects of cAMP microdomains on tumor growth, NSG mice were injected into the right flank with 5000 cells in 50 μL of 25% Matrigel Basement Membrane Matrix (Corning, #354248). Tumor growth was monitored every 3-4 days and the largest diameter was recorded. When tumors reached 1 cm in diameter, mice were randomized and one cohort was switched to doxycycline diet. When tumors in any cohort reached 2.5 cm in diameter, mice were euthanized, and tumor samples were collected for further analyses. Tumor growth data are presented as a fold change in size, normalized to tumor size on the day when mice were randomized and doxycycline was introduced into the diet. Upon euthanasia, tumors were removed, weighed, and samples were placed in formalin or flash frozen for subsequent analysis. Total number of mice per condition (regular chow/doxycycline chow): NLS-sAC 14/14, NES-sAC 9/9, mito-sAC 10/10, LacZ CTRL 5/5, 5SA-YAP 5/4, S397A-YAP 5/5, WT-YAP 5/5.

### Immunohistochemistry

Tumor sections were fixed in 10% formalin solution (Sigma, #HT501128) and paraffin embedded. IHC was performed on the Leica Bond RX using the Bond Polymer Refine Detection Kit (#DS9800). The sections were pre-treated using heat mediated antigen retrieval with EDTA pH9 (Leica Biosystem Epitope Retrieval Solution 2, #AR9640) for 20 minutes. The sections were then incubated with R21 (Anti-sAC antibody, CEP Biotech, Inc, 1:1200) or Ki67 (CST, #12202, 1:500). Leica BOND red (R21) or diaminobenzidine (Ki67) was used as chromogen then counterstained with hematoxylin. Slides were imaged on an Olympus light microscope with 10x objective and DP71 camera. R21 staining was quantitated manually by GD and nuclear staining was compared to depth of tumor invasion (Breslow), clinical outcome, and the percentage and nuclear positive tumor cells was assessed. Quantification of Ki67 positive nuclei was performed with Fiji/ImageJ using raw data images. Color deconvolution H/DAB was applied, followed by RenyiEntropy threshold, and watershed tool to count the Ki67 positive nuclei per field (on average ∼4000 nuclei/image from doxycycline free condition were detected). 2-5 fields of non-necrotic sections were imaged per tumor, five tumors/condition (except NLS- sAC/ regular chow, NES-sAC/doxycycline chow, and 5SA-YAP/doxycycline chow which were n=4). Data in graphs are presented as a mean fold change over corresponding doxycycline-free condition ± SEM. For the purpose of figure preparation, contrast and brightness was adjusted by applying exactly the same settings to images from all the groups.

### Cell culture and drug treatment

Generation of *Adcy10^-/-^* and *Adcy10 ^WT/^*^WT^ melanocytes is described elsewhere(Zhou et al., 2018). Mouse melanoma Mel 2-4 line (sAC^KO^) was derived from tumors formed by *Adcy10^-/-^* melanocytes injected subcutaneously into immunocompromised mice. Tumors were dissociated in trypsin immediately after removal from the mouse, placed in RPMI media containing 10% bovine serum and antibiotics, and allowed to adhere to plastic tissue culture plates. “SK-Mel” and Yummer1.7 melanoma cell lines were provided by Taha Merghoub (Memorial Sloan-Kettering Cancer Center, New York, NY), SCC12 was provided by Loraine Gudas (Weill Cornell Medical College, New York, NY), colon cancer lines (DLD1 and SW480) were provided by Lukas Edward Dow (Weill Cornell Medical College, New York, NY), pancreatic cancer lines (Hs766t, PANC1, MIAPaCa2) were provided by Lewis Cantley (Weill Cornell Medical College, New York, NY), prostate cancer cell line LNCaP was provided by Christopher Barbieri (Weill Cornell Medical College, New York, NY) and the melanoma M263 line was provided by Roger Lo (UCLA Medical Center). Mel 2-4, SK-Mel, M263 and prostate lines were maintained in RPMI, pancreatic and colon lines in DMEM while SCC12 and Yummer1.7 in DMEM/F12 medium each supplemented with 2 mM glutamine, 50 U/ ml of penicillin/ streptomycin, and 10% heat-inactivated fetal bovine serum. Additionally, SCC12 medium was supplemented with 400 ng/ml hydrocortisone, and Yummer1.7 medium with non-essential amino acids (Gibco, #11140-050). Cells were maintained at 37°C in 5% CO_2_.

Drugs were prepared according to manufacturers’ instructions and added directly to cell culture medium. Doxycycline hyclate (Sigma, #D9891) was used as indicated. IBMX (3- Isobutyl-1-methylxanthine; Sigma, #I7018) was used at 50 or 500 μM. PGE2 (Prostaglandin E2; Tocris, #2296) was used at 10 μM. Isoproterenol hydrochloride (Calbiochem, #420355) was used at 10 μM. LRE1 (gift from Lonny Levin and Jochen Buck) was used at 50 μM. KT5720 (Sigma, #K3761) was used at 10 μM. TRULI (gift from the Tri-I TDI) was used at 500 nM. Leptomycin B (Cayman Chemical Company, # 10004976) was used at 46 nM. Cycloheximide (Sigma, # 01810) was used at 20 μg/ml.

Bicarbonate stimulation of human melanoma cells was performed under ambient CO_2_ conditions at 37°C. Cells were cultured in pH 7.2 stabilized media deficient in bicarbonate for 6h, and then stimulated by replacing media with regular culture media containing bicarbonate.

### Plasmid DNA, primers and cloning

The following plasmids were obtained from Addgene: pCW57.1 (Addgene plasmid, #41393); pcDNA3-ICUE3 (Addgene plasmid, # 61622)(DiPilato and Zhang, 2009). GFP-PKI nls (Addgene plasmid, #118301)(Billiard et al., 2001); pLL3.7-EF-EYFP-YAP1_WT-PolyA (Addgene plasmid, #112284)(Ege et al., 2018); pQCXIH-Myc-YAP (Addgene plasmid, #33091)(Zhao et al., 2007); pQCXIH-Myc-YAP-5SA (Addgene plasmid, #33093)(Zhao et al., 2007); pQCXIH-Flag-YAP-S127A (Addgene pasmid, # 33092)(Zhao et al., 2007); and pQCXIH-Flag-YAP-S381A (Addgene plasmid, #33068)(Zhao et al., 2010). Of note, YAP S381 in mice corresponds to residue S397 in humans; all reference to phosphorylation at this site was described as S397.

EYFP-YAP-S397A plasmid was created from pLL3.7-EF-EYFP-YAP1_WT-PolyA by site directed mutagenesis, and confirmed by sequencing.

LATS1 and 2 knock-down was achieved using GIPZ lentiviral plasmids specific for mouse LATS1 or 2 mRNA (Dharmacon #RMM4532-E-EG50523, and #RMM4532-EG16798).

sACt(Buck et al., 1999) was tagged with either two NLS or NES sequences(Dang and Lee, 1988; Fu et al., 2012; Wen et al., 1995). Mito-sAC(Acin-Perez et al., 2009a) was provided by Giovanni Manfredi (Weill Cornell Medical College, New York, NY). Gateway cloning system (Thermo) was used to clone sAC into the pCW57.1 plasmid. LacZ was shuttled into pCW57.1 from the pLenti6.3_V5-GW_lacZ plasmid.

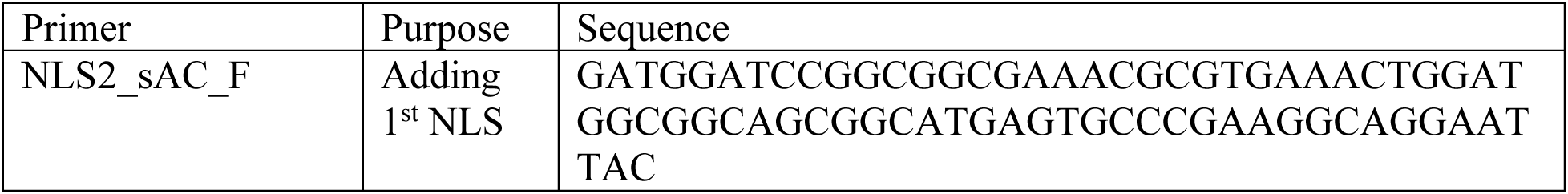
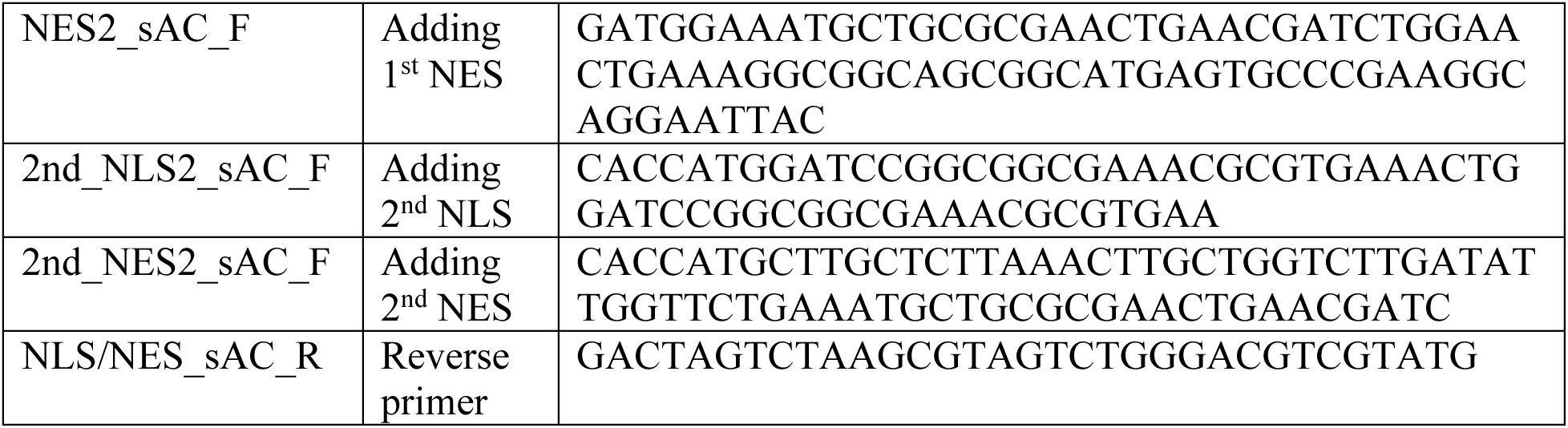

### Intracellular cAMP ELISA

Cells were seeded in 24-well plate at 2 x 10^5^ cells per well. Doxycycline was added at 1μg/ml for the time indicated. 500 μM 3-Isobutyl-1-methylxanthine (IBMX, Sigma, #I7018) was supplemented to the media for the indicated time. Adherent cells were lysed directly in 200 μl of 0.1 M HCl, and the intracellular cAMP content was determined using the cAMP complete ELISA kit (ENZO Life Sciences, # ADI-901-066). Data is represented either as a fold change in cAMP level over untreated cells, or pg/ml cAMP accumulated per 10^5^ cells.

### Lentivirus generation and clone selection

293T cells were transfected with pMD2.G enveloping plasmid, psPAX.2 packaging plasmid, and either plasmids encoding microdomain-targeted sAC (pCW57.1 constructs), or YAP variants (pQCXIH constructs). Medium containing viral particles was collected approximately 60h later, passed through a 0.45 µm filter and used immediately. For infection, lentivirus was added with polybrene (Sigma, # H9268) at 6 µg/ml. The next day cell culture medium was replaced, and drug selection was started 24h later. Puromycin dihydrochloride (Sigma, #P8833) was used to select sAC expressing cells at 1-7.5 μg/ml depending on the cell line. Hygromycin B (Sigma, #H3274) was used at 200 μg/ml to select cells expressing YAP variants.

### Immunocytochemistry and fluorescent light microscopy

Cells were seeded on glass coverslips and grown as a monolayer. They were fixed with 4% paraformaldehyde (Electron Microscopy Sciences, #15710) for 10 min, permeabilized in 0.5% Triton X-100 (Sigma, #X100) in PBS for 5 min, and blocked in 3% bovine serum albumin (Proliant Biologicals, #68100) for one hour. Immunolabeling with primary and secondary antibodies was conducted for one hour at room temperature. Primary antibodies targeted: HA-Tag (C29F4; CST, #3724; dilution 1:500), YAP (D8H1X; CST, #14074; dilution 1:100), and cytochrome C (2CYTC-199; Santa Cruz, #sc-81752; dilution 1:50). Secondary antibodies used were donkey anti-mouse and donkey anti-rabbit conjugated to Alexa Fluor 488 or Alexa Fluor 647 (Thermo, #A-21206, #A-31571, #A-31573) and diluted 1:300. Coverslips were mounted on glass microscopy slides with ProLong Gold Antifade Mountant with DAPI (Thermo, #P36931). Immunostained cells were imaged on a Zeiss LSM 880 Inverted laser scanning microscope (Zeiss, Oberkochen, Germany) using a Plan-Apochromat 40x/1.3 Oil DIC M27 immersion lens. Images were acquired using ZenBlack operating software. Any quantitation performed, was done using raw data images. For the purpose of figure preparation, contrast and brightness was adjusted by applying exactly the same settings to images from control and experimental groups.

### YAP nuclear localization analysis

Microscopy images were acquired as described above. Quantitative analyses were performed using ImageJ 2.0 (Wayne Rasband, NIH, USA). Nuclei were identified by DAPI stain. To define regions of interest, the YAP microscopy image threshold was set to pixel intensity greater than 10 and converted to binary image. Erosion, followed by dilation operation, was performed to remove isolated pixels. Such regions of interest were applied to raw YAP microscopy images and the total pixel intensity, corresponding to the total YAP signal was calculated per cell. The nuclear region was defined by DAPI stain, and used to calculate the nuclear YAP signal on a per cell basis. Cytoplasmic YAP signal was determined by subtracting nuclear YAP signal from the total YAP signal. Nuclear to cytoplasmic ration (N/C) was calculated per cell. To study effects of PKI on YAP cellular localization, cells were transfected with NLS-PKI-GFP expressing plasmid (Addgene, #118301), and used for experiments 24h later. NLS-PKI expressing cells were identified by GFP fluorescence. Data was normalized to untreated control cells and presented as mean with SEM. Each experimental condition was repeated 3-5 times.

### YAP nuclear export analysis

Fluorescent recovery after photobleaching (FRAP) was performed on a Zeiss LSM 880 Inverted laser scanning microscope (Zeiss, Oberkochen, Germany) fitted with temperature-controlled and CO_2_ chamber for live cell work using a Plan-Apochromat 40x/1.3 Oil DIC M27 immersion lens. Images were acquired using ZenBlack operating software. EYFP-YAP expressing cells were seeded in 35 mm glass bottom dishes (MatTek). After 24 h incubation with or without doxycycline, nuclear region was bleached with 405 nm laser wavelength, to reduce nuclear EYFP-YAP signal by at least 70%. Recovery of EYFP-YAP signal inside the nucleus was measured every 5 seconds over indicated time. Mean fluorescent intensity data was normalized by setting the initial fluorescence signal (pre-bleach) to 100%, and the signal immediately after photo-bleaching (timepoint 0) to 0%. Normalized recovery of EYFP-YAP signal inside the nucleus is presented from timepoint 0s. EYFP-YAP fluorescence in adjacent, non-bleached nuclei remained constant.

### Intracellular cAMP measurement in live cells

Cells were transfected with the cAMP FRET probe ICUE3(DiPilato and Zhang, 2009; Sample et al., 2012b). Briefly, cells were seeded in a 24-well µ-Plate (iBiDi, #82406) and grown overnight either with or without 1 µg/ml doxycycline. Next day, cells were washed twice and maintained in Hank’s balanced salt solution in a CO_2_-independent incubator. After 10 min equilibration time, they were treated with 15 mM NaHCO_3_. Images were acquired on a Zeiss LSM 880 Inverted laser scanning microscope (Zeiss, Oberkochen, Germany) fitted with temperature-controlled chamber for live cell work using a Plan-Apochromat 40x/1.3 Oil DIC M27 immersion lens. Images were acquired every 30 s using ZenBlack operating software. Dual emission ratios were obtained with 405 nm wavelength excitation, and emission filters cycled between 475 / 40 nm for cyan fluorescent protein and 535 / 25 nm for yellow fluorescent protein present in ICUE3. Images were analyzed in ImageJ 2.0 (Wayne Rasband, NIH, USA). Background correction was performed by subtracting autofluorescence of regions without cells from the emission of cells expressing ICUE3. Intensity of cyan and yellow fluorescent protein was measured in both cytoplasmic and nuclear regions within each cell and cyan to yellow ratio was calculated. The ratio change over the bicarbonate stimulation was then normalized to the ratio of image acquired immediately after adding bicarbonate (set to 1).

### Immunoblotting

Primary antibodies used for Western Blot analysis: HA-Tag (C29F4; CST, #3724), YAP (D8H1X; CST, #14074), Phospho-Ser397 YAP (D1E7Y; CST, #13619), Phospho-Ser127 YAP (D9W2I; CST, #13008), Phospho-Ser109 YAP (CST, #46931), Lats1 (C66B5; CST, #3477), Lats1 (CST, #9153), Phospho-Thr1079 Lats1 (D57D3; CST, #8654), Phospho-Ser909 Lats1 (CST, #9157), TAZ (D3I6D; CST, #70148), Phospho-Ser89 TAZ (E1X9C; CST, #59971), GAPDH (14C10, CST, #2118), H3 (Abcam, #ab1791), and Beta-tubulin (TUB2.1, Sigma, #T4026). All antibodies were used at the manufacturer’s suggested dilutions. Secondary antibodies: HRP linked anti-rabbit IgG (CST, 7074), and HRP linked anti-mouse IgG (Amersham, #NXA931) were used at the manufacturer’s recommended dilutions. Cultured cells were lysed directly in SDS-PAGE sample loading buffer (50 mM Tris-Cl, pH 6.8; 2% SDS, 6% (v/v) glycerol; 0.01% (w/v) bromophenol blue) containing 5% 2-mercaptoethanol, heated at 95°C for 5 min, electrophoresed in 8 or 10% Tris gels, and transferred to nitrocellulose membranes (GE Life Sciences, #10600002). The membranes were blocked in TBST (Tris-buffered saline, 0.1% Tween 20) containing 5% Bovine Serum Albumin. Antibodies were diluted in blocking buffer. Incubation with primary antibodies was performed at 4°C overnight, while incubation with HRP-conjugated secondary antibodies was performed at room temperature for one hour. Protein bands were detected with SuperSignal West Femto enhanced chemiluminescence substrate (Thermo, #34096) or HyGlo Quick Spray (Denville, #E2410). When necessary, the membranes were reprobed after incubation in Restore Western Blot stripping buffer (Thermo, #21059). GAPDH or Beta-tubulin antibodies were used as loading controls. Tumor samples were homogenized in RIPA buffer (Thermo, #89901) followed by protein concentration measurement using DC Protein Assay (Bio-Rad, #5000112). Tumor lysates were immunoblotted as above. Membranes were stripped with Western Blot Strip-It Buffer (Advansta, # R-03722-D50). Quantitation of Western blot band volumes was performed in Image Lab 6.1 (Bio-Rad Laboratories) using non-saturated raw data images. For the purpose of figure preparation, black color balance was adjusted to 10% across all the Western images shown in the paper.

### Nuclear fractionation

Fractionation of intact nuclei from cytoplasm was performed using Nuclei Isolation Kit: Nuclei EZ Prep (Sigma, #NUC101-1KT) following manufacturer’s instructions. For Western blot analysis equal cell equivalents were loaded for nuclear and cytoplasmic fractions.

### In vitro cancer cell growth

#### Matrigel assay

In vitro cancer cell growth in 3D matrix was performed as previously described(Pauli et al., 2017) with modifications. 96-well plates with white walls and clear bottom (Corning, #3610) were coated with 40 μl of 3 mg/ml Matrigel matrix solution (Corning, #356231). Cell suspension was mixed with Matrigel (1:2) and 1000 cells were seeded per well. Wells were topped with culture media with or without doxycycline, and kept in a cell incubator (37°C, 5% CO_2_) for 7-14 days until organoids formed. Media was replenished every 2-3 days. Viability of tumor organoids was assessed by CellTiter-Glo® 2.0 assay (Promega, #G9242) according to manufacturer’s instructions. Luminescence was measured using a SpectraMax plate reader.

#### Clonogenic assay

3000-5000 cells/well were seeded in 6 well plates and incubated at 37°C in CO_2_ incubator. The next day doxycycline was added to wells. Media was changed every 2-3 days. After 11-14 days, the plates were washed with phosphate buffered saline (PBS), fixed with 10% formalin for 40 minutes, stained with 0.01% crystal violet diluted with dH_2_O for 1 hour, rinsed with dH_2_O, and allowed to air dry overnight. Colony formation was assessed by photography and by dissolving the crystal violet stained cells with 10% acetic acid (Spectrum Chemical, #AC110) solution diluted in dH_2_O for 30 minutes. Crystal violet absorbance was measured at 590nm using a SpectraMax plate reader.

### Genomic data analysis

#### RNA-seq

Total RNA was isolated using the RNeasy mini kit (Qiagen, # 74104). Following RNA isolation, total RNA integrity was checked using a 2100 Bioanalyzer (Agilent Technologies, Santa Clara, CA). RNA concentrations were measured using the NanoDrop system (Thermo Fisher Scientific, Inc., Waltham, MA). Preparation of RNA sample library and RNA-seq were performed by the Genomics Core Laboratory at Weill Cornell Medicine. Messenger RNA was prepared using TruSeq Stranded mRNA Sample Library Preparation kit (Illumina, San Diego, CA, # 20020595), according to the manufacturer’s instructions. The normalized cDNA libraries were pooled and sequenced on Illumina HiSeq4000 sequencer with pair-end 50 cycles.

Transcript abundances were quantified using Salmon version 1.2.0(Patro et al., 2017) by selective alignment with a decoy-aware transcriptome index of Gencode Release M23 (GRCm38.p6, https://www.gencodegenes.org/mouse/release_M23.html). Transcript abundance estimates from Salmon were imported into R and summarized to annotated gene counts using the BioConductor package TXImeta(Love et al., 2020). Differential expression was computed using the BioConductor package DESeq2(Love et al., 2014). Gene set enrichment analysis (GSEA) was performed using the genome-wide Log_2_ fold-changes for each microdomain. Gene sets with positive enrichment were reported as up-regulated, and those with negative enrichment were reported as down-regulated, and significance was determined using a 1% FDR.

Number of tumor samples used for RNA-seq analysis was n=2 per cell type (NLS-sAC, NES-sAC, mito-sAC) per condition (regular chow vs doxycycline chow) for a total of n=12. RNA-seq on cells treated in vitro was performed on three NLS-sAC clones, with (48h) or without doxycycline, in parallel with two LacZ control lines, with (48h) or without doxycycline, for a total of n=10.

Sequencing data can be found at: https://www.ncbi.nlm.nih.gov/geo/query/acc.cgi?acc=GSE154877.

#### ATAC-seq

ATAC-seq library preparation and sequencing was performed at the Epigenomics Core at Weill Cornell Medicine by using the OMNI-ATAC-seq method described by Corces(Corces et al., 2017). Briefly, 50,000 cells at ∼70% viability were spun down and incubated 3 min at 4°C in 25µl of a detergent buffer containing 0.2% Igepal CA-630 (Sigma, #I8896), 0.2% Tween 20 (Sigma, # P9416) and 0.02% Digitonin (Promega Corporation, #G9441). Nuclei were centrifuged at 500 g for 10 min and immediately resuspended in 25 µl of buffer containing 2.5 µl of Tn5 transposase (Illumina, #15027865) for a 30 min incubation at 37°C. Fragments generated by the Tn5 transposase were purified using the DNA Clean and Concentrate kit (Zymo Research, #D4014). Uniquely indexed libraries were obtained by amplification of the purified fragments with indexed primers using 10 cycles of PCR (5 min x 72 °C, 5 cycles each 10 sec x 98 °C, 30 sec x 63 °C, 1 min x 72 °C). Resulting libraries were subjected to a two-sided size clean up using SPRI beads (Beckman Coulter, Brea, CA) to obtain sizes between 200-1000bp, and pooled for sequencing. The pool was clustered at 9 pM on a pair end read flow cell and sequenced for 50 cycles on an Illumina HiSeq 2500 to obtain ∼40M reads per sample. Primary processing of sequencing images was done using Illumina’s Real Time Analysis software (RTA) as suggested by the Illumina. CASAVA 2.17 software was used to perform image capture, base calling and demultiplexing of the raw reads. Paired-end 50 base pair ATAC-seq reads were trimmed to remove adapter sequences using NGmerge with options “-u 41 -a”(Gaspar, 2018). Trimmed read pairs were aligned to version 38 of the mouse reference genome (GRCm38) using bowtie2 with the following options: “-X2000 --local --mm -k 4“. Aligned reads were filtered and sorted to exclude reads mapping to mitochondrial DNA and black-listed regions, and duplicate read pairs were removed using the “MarkDuplicates” program in picard tools (http://broadinstitute.github.io/picard), resulting in a final aligned, sorted, and filtered BAM file that was used for all subsequent analysis.

ATAC-seq peaks were called from Tn-5 corrected insertions using MACS2 callpeak with option “-g hs --nomodel --shift -75 --extsize 150 --keep-dup all --call-summits”. A chromatin accessibility atlas containing 500 bp disjoint genomic intervals (DNA elements) was constructed from called peak summits across all primary cells using an iterative peak-ranking method as previously described(Corces et al., 2018).To quantify accessibility across samples, the number of single-base Tn5-corrected insertions that fell within each 500 bp DNA element was counted from ATAC-seq bam files using the command “pyatac counts” in the nucleoATAC package(Schep et al., 2015). Differential accessibility between conditions was computed using DESeq2. Normalization factors included in the call to DESeq2 were computed by quantile normalization with GC sequence content bias correction using the EDAseq BioConductor package. Chromatin accessibility Log_2_ fold-changes were computed and shrunken using the function “lfcShrink” with option “type=ape” in the DESeq2 R package(Zhu et al., 2019). Differentially accessible DNA elements were identified at FDR of 10%. All gene-based annotation was performed using Gencode Release M23 (GRCm38.p6, https://www.gencodegenes.org/mouse/release_M23.html). DNA elements were assigned to genes according to the nearest TSS of a protein coding gene.

Annotated transcription factor motifs were identified in genomic DNA spanning the 500bp DNA elements using the R package motifMatchR version 1.8.0 (https://bioconductor.org/packages/release/bioc/html/motifmatchr.html) with the options “p.cutoff=1e-6”. A filtered and curated collection of mouse transcription factor motifs was obtained from chromVARmotifs (“mouse_pwms_v2”, https://github.com/GreenleafLab/chromVARmotifs)(Schep et al., 2017). Transcription factor motifs were filtered to include those in which the corresponding transcription factor was expressed with transcript abundance > 1 TPM in at least one replicate experiment (see RNA-seq methods). To calculate change in accessibility remodeling, we performed a linear regression of the shrunken Log_2_ fold-changes on each TF motif independently, resulting in 1 regression model for each expressed TFs. Hypothesis testing of estimated coefficients was performed by Wald-test with a heteroskedasticity-consistent robust covariance estimator, using the “sandwhich” package in R. TF coefficient estimate p-values were corrected for multiple-hypothesis testing and the top significant TFs were reported (FDR q-value<1e-5).

ATAC-seq on cells treated in vitro was performed on three NLS-sAC clones, with (48h) or without doxycycline, in parallel with three LacZ control lines, with (48h) or without doxycycline, for a total of n=12.

Sequencing data can be found at: https://www.ncbi.nlm.nih.gov/geo/query/acc.cgi?acc=GSE154877.

#### Comparison of hyperactive YAP (5SA allele) and NLS-sSAC associated RNA expression changes

FASTQ files were acquired from the authors of Zhang et al. (2020)(Zhang et al., 2020b), each containing single-end RNA-seq reads from one of three biological replicates of MeWo cells expressing a DOX inducible vector control or hyperactive YAP, either untreated or treated with DOX. Quantification of gene expression was performed using Salmon(Patro et al., 2017) (version 1.4.0) using the seqBias flag. Sequencing reads were automatically inferred to be forward stranded by Salmon. The Salmon index was generated using kmers of length 31 from mature transcript sequences created from Ensembl gene set annotations (release 104) and the GRCh38 primary assembly using rtracklayer (version 1.50.0)(Lawrence et al., 2009), GenomicRanges (version 1.42.0)(Lawrence et al., 2013), and Biostrings (version 2.58.0) (Pages et al., 2021) in R. The whole genome was used as a decoy for selective alignment.

To measure hyperactive YAP associated changes in gene expression, we used DESeq2 (version 1.30.1)(Love et al., 2014). Transcript level counts from Salmon were summarized to gene counts using tximport (version 1.18.0)(Soneson et al., 2015) with the countsFromAbundance argument set to ‘lengthScaledTPM’. We modeled gene counts in MeWo cells as a linear function of YAP status (vector control vs YAP-5SA), DOX treatment, and an interaction term between YAP status and DOX treatment. The interaction coefficient corresponds to DOX-induced gene expression changes (in log base 2) in YAP-5SA cells relative to vector control.

To compare hyperactive YAP associated gene expression changes in MeWo cells to NLS-sAC associated changes in mouse cells, we identified high confidence orthologues with 1- to-1 mapping between humans and mice in the Ensembl database. We classified these genes as up- or down-regulated in each respective RNA-seq experiment (i.e., MeWo YAP-5SA vs control experiment, in vivo and in vitro NLS-sAC vs control experiments), then applied Fisher’s exact test to a 2×2 contingency table to assess the concordance of observed changes between experiments. The expected counts per contingency table cell were computed assuming the proportion of up- and down-regulated genes in one experiment is independent from the other experiments, e.g., to get the expected number of genes downregulated in both experiments being compared, we simply multiplied the proportion of downregulated genes in the first experiment by the proportion of downregulated genes in the second experiment by the total number of genes.

#### Inference and comparison of YAP and NLS-sAC activities in human melanoma cell lines

Melanoma cell line data from the Cancer Cell Line Encyclopedia (CCLE)(Ghandi et al., 2019) were acquired from the DepMap data portal(Tsherniak et al., 2017) (release 20Q1; file names: sample_info.csv, CCLE_RNAseq_reads.csv, CCLE_mutations.csv). Mutation data were used to exclude genetically related melanoma cell lines from the analysis. For each pair of cells that shared a conservative percentage of at least 10% of the union of their called mutations, we first prioritized the one whose DepMap_ID was listed in the sample_info file, then the one which had corresponding RNA-seq data, then the one which lacked additional information (additional_info) in the sample_info file, a field potentially indicating the cell line was derived from a parental cell line, such as through genetic modification or drug selection. Lastly, we prioritized the cell line with the most mutation calls, and in case of ties, we prioritized the cell line whose DepMap_ID ranked first alphanumerically when IDs were sorted from smallest to largest. We retained 45 of the 62 original melanoma cell lines with RNA-seq data after filtering. The retained cells line names are listed at the bottom of figure 6c.

RNA-seq counts for the 45 melanoma cell lines were loaded into R. Genes with low counts were flagged and removed using the edgeR package’s (version 3.32.1) filterByExpr function and default parameters(McCarthy et al., 2012). Read counts were normalized for differences in sequencing depth and subjected to variance stabilizing transformation using DESeq2.

Normalized transformed gene expression values were used to estimate YAP and NLS- sAC activity per cell line using the Gene Set Variation Analysis (GSVA) R package (version 1.38.2)(Hanzelmann et al., 2013). Briefly, GSVA standardizes expression values for each gene, then uses a Kolmogorov-Smirnov-like rank statistic to infer if genes in a provided gene set are relatively highly expressed, lowly expressed or not differentially expressed from genes outside the set, producing a per-sample continuous score greater than, less than, or equal to zero, respectively.

We provided GSVA with four gene sets. We used genes downregulated upon NLS-sAC induction in our in vitro experiments as markers of NLS-sAC activity. In one set, we included genes with a log fold-change < 0 and FDR < 10% (NLS-sAC signature 1; n = 460 genes), and in another set we included genes ranked in the bottom 1% based on the Wald statistic from DESeq2 (NLS-sAC signature 2; n = 100 genes). We used genes upregulated upon YAP-5SA induction in MeWo cells as markers of YAP activity. In one set, we included genes with a log fold-change > 0 and FDR < 10% (YAP signature 1; n = 470 genes), and in another set we included genes ranked in the top 1% based on the Wald statistic from DESeq 2 (YAP signature 2; n = 126 genes). GSVA was run using default parameters (method = “gsva”, kcdf = “Gaussian”, abs.ranking = FALSE, mx.diff = TRUE). We transformed NLS-sAC scores from GSVA to activity scores by multiplying them with -1.

To validate the robustness of our protein activity estimates, we complemented GSVA with an alternative approach. We first derived NLS-sAC and YAP activity markers by ranking genes based on their absolute Wald statistic then selecting the top 1% of genes and setting aside their log fold-changes, estimated by DESeq2 (in vitro NLS-sac up n = 45 genes, in vitro NLS- sAC down n = 55 genes, MeWo YAP-5SA up n = 107 genes, MeWo YAP-5SA down n = 19 genes). We centered the variance stabilized expression values of these genes around their mean in CCLE melanoma cell lines. We computed the log2 odds-ratio between the sign of the DESeq2 log fold-changes with the sign of the centered expression values per sample. We used this log odds-ratio as an indicator for protein activity per cell line, where positive values indicated higher activity and negative values indicated lower activity (NLS-sAC signature 3 and YAP signature 3).

Clustering of melanoma cell lines in figure 6c was derived using the consensus clustering, implemented in the ConsensusClusterPlus R package (version 1.54.0)(Wilkerson and Hayes, 2010). We applied ConsensusClusterPlus on variance stabilized gene expression values for 10,000 clustering iterations, at each iteration sampling 80% of cell lines, clustering them using Pearson’s distance and Ward’s linkage (ward.D2), then classifying them into two groups by cutting the resulting dendrogram at its highest banchpoint. The resulting “consensus matrix”, which contains the proportion of times each pair of cells were grouped together, was clustered using Ward’s linkage then used to classify cells into two final groups. The genes were similarly clustered by resampling subsets of genes for 100 iterations.

### RT-PCR

Cell pellets were collected in RNAlater (Sigma Aldrich, #R0901) and stored at -20° C. Samples were homogenized using Qiashredder columns (Qiagen, #79656) and RNA was isolated according to specifications of the RNeasy plus mini kit (Qiagen, #74136). cDNA was made using the high-capacity RNA-to-cDNA kit (Thermo Fischer, #4387406). The applied biosystems power SYBR green PCR master mix (Thermo Fischer, #4368706) was used for qPCR using the QuantStudio 6 real-time PCR instrument (Thermo Fischer). Delta delta CT analysis was performed to determine relative mRNA expression amongst samples normalized to *GapDH*.

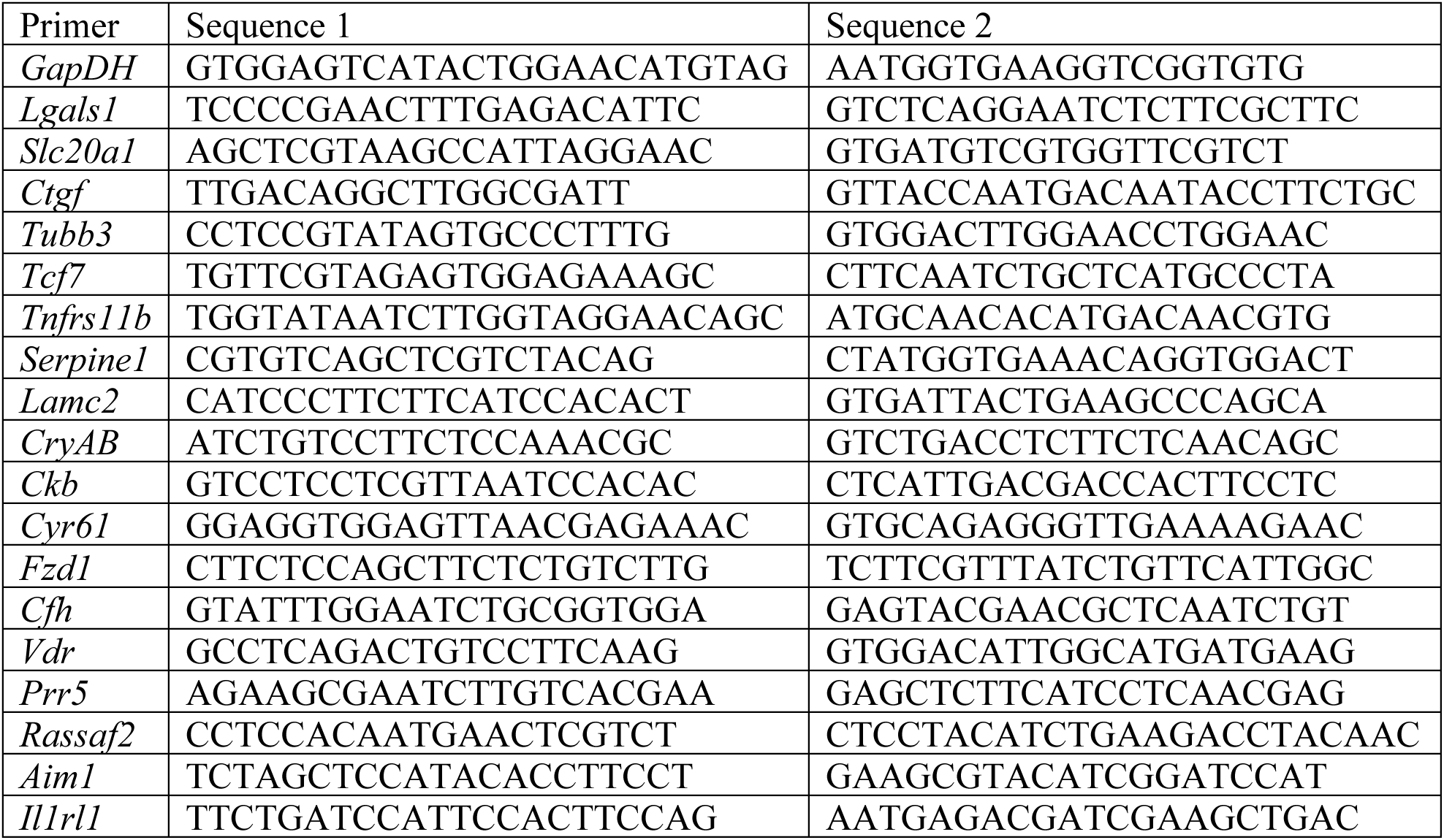
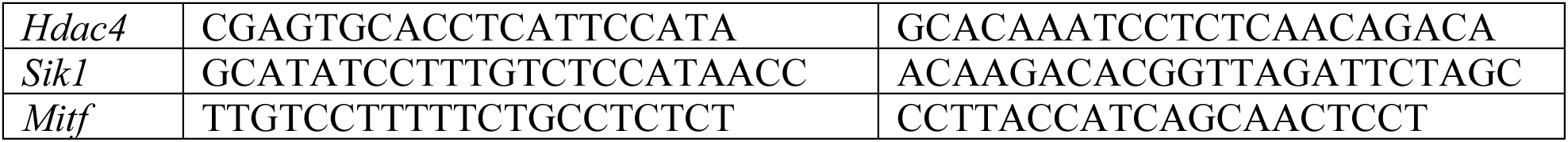

### Statistical analysis

All statistical analyses were performed using GraphPad Prism 8.0 (GraphPad Software). Experiments were performed a minimum of three times, unless indicated otherwise. Comparison of means was performed using either an unpaired, two-tailed *t* test for two groups, or an ANOVA with Sidak correction for multiple comparisons for groups of three or more. A two-sided Fisher’s exact test was performed on the contingency data presented for nuclear sAC staining in melanoma primary biopsy samples with established metastases. Wilson-Brown was performed to calculate sensitivity and specificity.

Binomial logistic regression was used to assess whether nuclear sAC positive melanoma (defined as >10% cells with nuclear staining by IHC) differed by Breslow thickness (Figure 1F). The response variable was the number of nuclear sAC positive samples and total samples, and the explanatory variable was Breslow thickness. Significance testing was performed using likelihood ratio test. Shaded regions represent the 95% confidence interval for percentage of nuclear sAC positive observations at each level of Breslow thickness.

## Supplemental Information

Figure S 1-10 with legends.

Supplemental Spreadsheets

## Acknowledgments

We thank members of the Zippin lab for critical reading of the manuscript and Paul Christos (Weill Cornell Medical College) for independent analysis of the statistics of the manuscript. We thank Dr. Jedd Wolchok for support for this work. We thank the following people for providing cell lines for this work: SCC12 were provided by Dr. Loraine Gudas (Weill Cornell Medical College, New York, NY), colon cancer lines (DLD1 and SW480) were provided by Dr. Lukas Edward Dow (Weill Cornell Medical College, New York, NY), pancreatic cancer lines (Hs766t, PANC1, MIAPaCa2) were provided by Dr. Lewis Cantley (Weill Cornell Medical College, New York, NY), prostate cancer cell line LNCaP was provided by Dr. Christopher Barbieri (Weill Cornell Medical College, New York, NY) and the melanoma M263 line was provided by Dr. Roger Lo (UCLA Medical Center). We thank Dr. Lukas Edward Dow for providing shRNA constructs against YAP and TAZ. We thank Peter T. Meinke of the Tri-Institutional Therapeutics Discovery Institute for providing TRULI inhibitor. We thank Kieran Harvey (Peter MacCallum Cancer Centre) for YAP 5SA MeWo RNAseq datasets.

