## Supplementary Figures S1-S10 for "A nuclear cAMP microdomain suppresses tumor growth by Hippo pathway inactivation"

**This PDF file includes:**

Figures S1 to S10

### Supplementary Figures

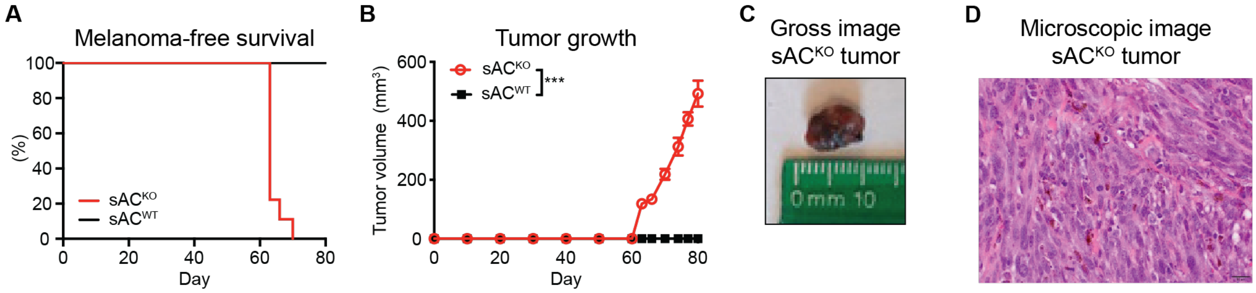


**Figure S1. sAC$^{KO}$ melanocytes form tumors more readily in mice.**
A) Melanoma-free survival in immunocompromised mice following subcutaneous injection with sAC knock-out (sAC$^{KO}$; red) or sAC wild type (sAC$^{WT}$; black) immortalized melanocytes (n=9 per cohort).
B) Tumor formation of sAC$^{KO}$ or sAC$^{WT}$ melanocytes injected subcutaneously into immunocompromised mice (n=9 per cohort; error bars, SEM; ***, $P \leq 0.001$).
C) Gross image of a sAC$^{KO}$ tumor.
D) Microscopic image of sAC$^{KO}$ tumor showing characteristics of invasive neoplasm composed of spindle cells; numerous cells display brown granules consistent with melanin depositions.

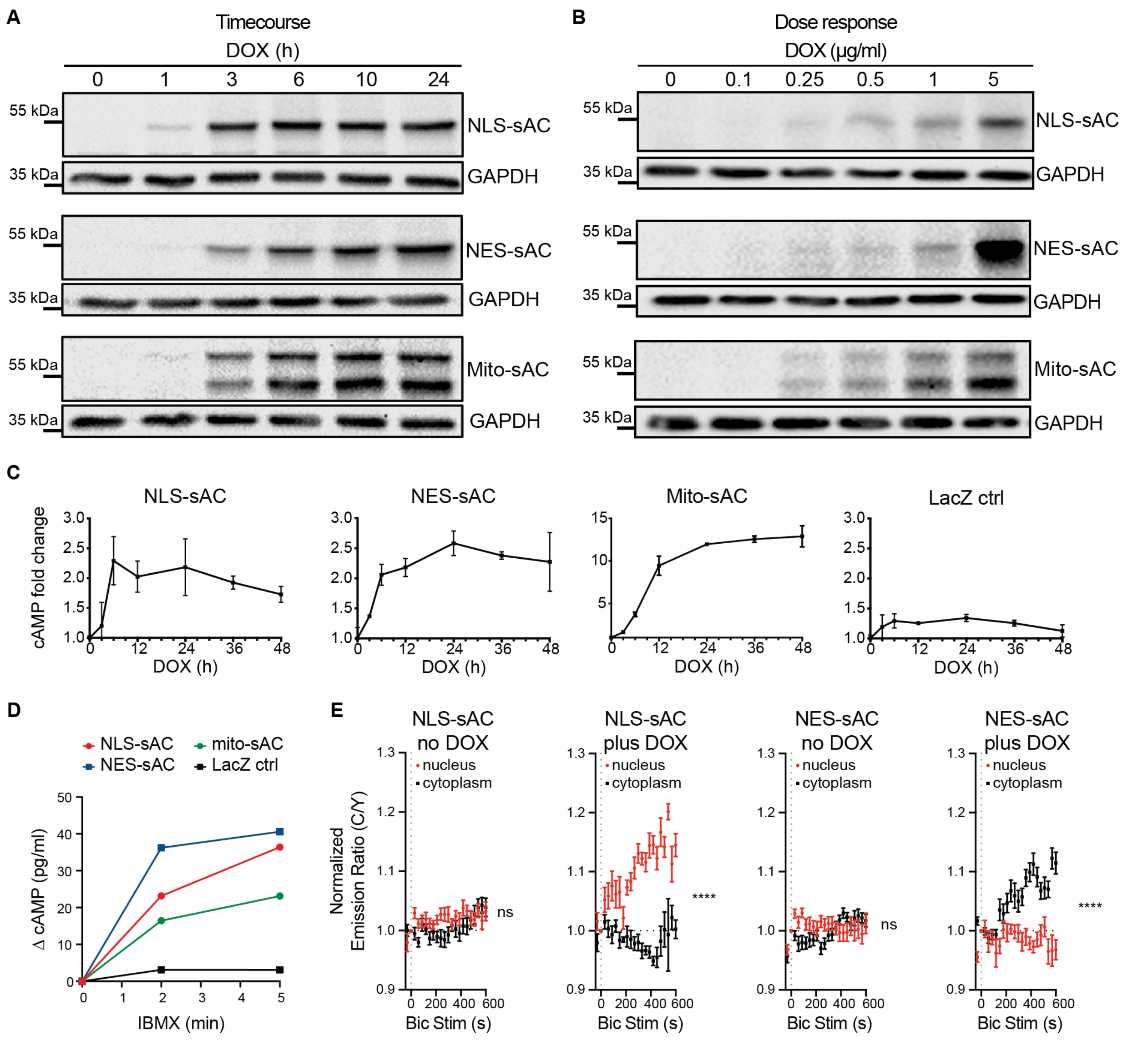


**Figure S2. Development of the sAC tool box for cAMP microdomain studies.**
A) Western blot analysis of doxycycline (DOX)-induced sAC expression over indicated times as determined by anti-HA antibody immunolabeling. Note the doublet in Mito-sAC which is a result of post-translational cleavage upon mitochondrial import.
B) Western Blot analysis of microdomain targeted sAC expression in response to increasing doses of doxycycline (DOX).
C) New cAMP equilibrium established in melanoma cells after doxycycline (DOX) induced sAC microdomain expression over indicated time as determined by the whole cell cAMP measured by ELISA. Error bars, SD.
D) Whole cell cAMP measured by ELISA from cells cultured for 24h in the presence of doxycycline, followed by PDE inhibition by IBMX (3-isobutyl-1-methylxanthine, 500 µM) for indicated times.
E) FRET measurements for cAMP probe ICUE3 in response to 15 mM $NaHCO_3$ stimulation in mouse melanoma cell lines. NLS- or NES-sAC expression was induced by 24h doxycycline treatment (plus DOX) prior to $NaHCO_3$ stimulation, and changes in CFP/YFP ratio,

corresponding to intracellular cAMP level, were measured simultaneously in both nuclear (red) and cytoplasmic (black) regions. Data normalized to timepoint zero defined as an image acquired immediately after adding $NaHCO_3$. NLS-sAC n=3; no DOX (19 cells), plus DOX (15 cells). NES-sAC n=2; no DOX (18 cells), plus DOX (19 cells). Mixed effect ANOVA (ns, $P > 0.05$; ****, $P \leq 0.0001$). Error bars, SEM.

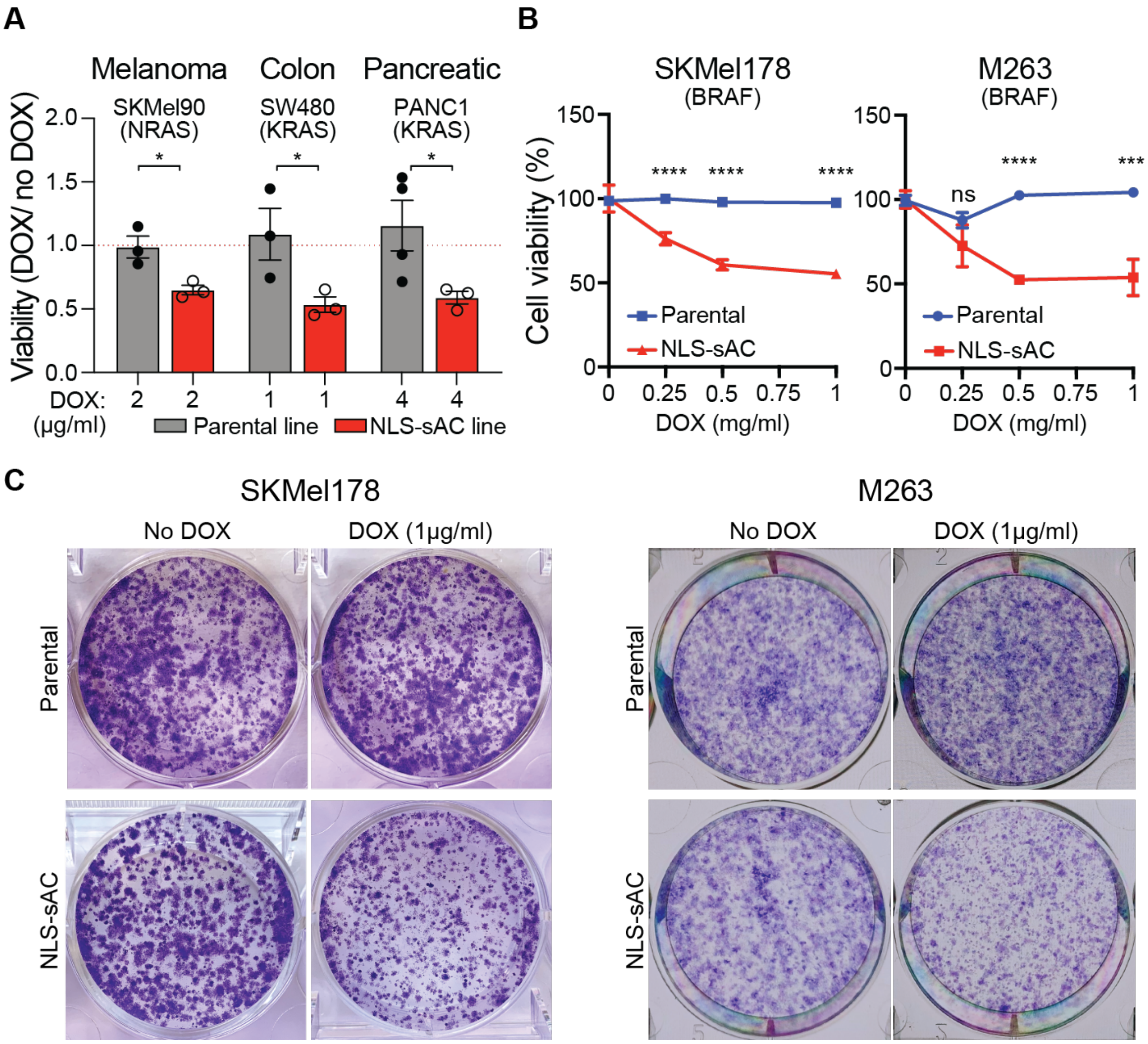


**Figure S3. In vitro growth inhibition of human cancers by nuclear cAMP.**

A) Growth inhibition of human cancer lines by nuclear cAMP in vitro in Matrigel, assessed by luminescence-based measurement of ATP content. Error bars, SEM; Student's t-test; DOX, doxycycline. n=3.

B) Clonogenic assay assessment of BRAF V600E melanoma cell line viability in response to induction of nuclear cAMP by doxycycline (DOX) as measured by absorbance of solubilized crystal violet normalized to no doxycycline. Error bars, SEM; ANOVA.

C) Image examples of the wells from the clonogenic assay in (B) following crystal violet staining.

(ns, $P > 0.05$; *, $P \leq 0.05$; ***, $P \leq 0.001$; ****, $P \leq 0.0001$).

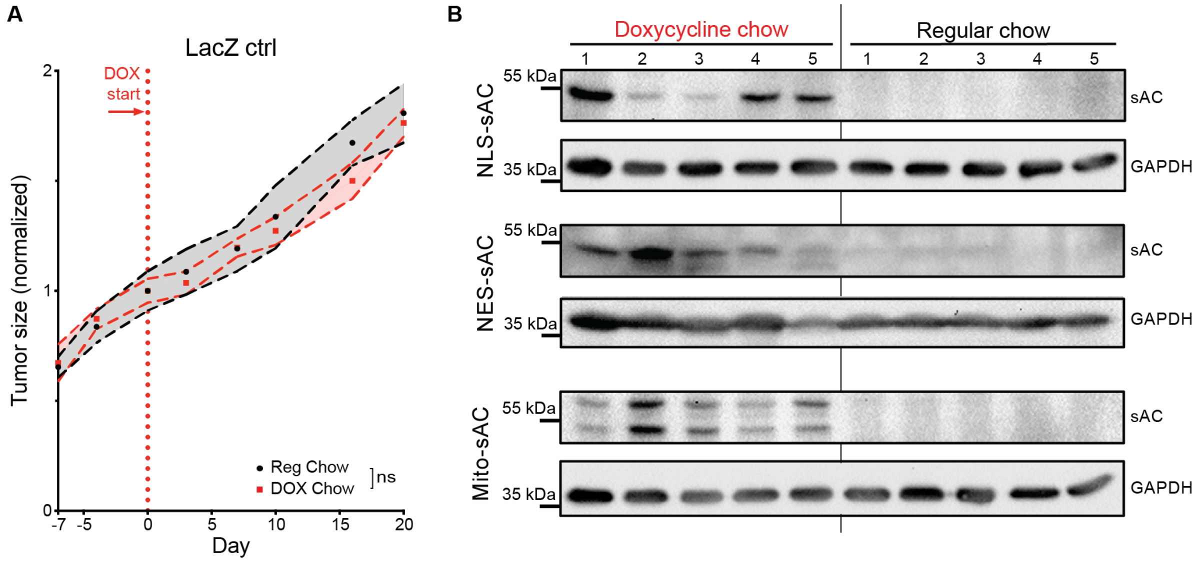


**Figure S4. Examination of cAMP microdomains in vivo.**
A) Tumor growth of LacZ control line in immunocompromised mice. Switch to doxycycline containing chow (DOX, red) is indicated by the arrow and red line. Data normalized to day 0 (day when mice changed to DOX chow). Reg, regular chow, gray. Error bars, SEM; two-way ANOVA; n=5 per cohort; ns, $P > 0.05$
B) Examples of Western blot analysis of tumor lysates showing expression of sAC microdomains upon switch to doxycycline chow as detected by anti-HA antibody.

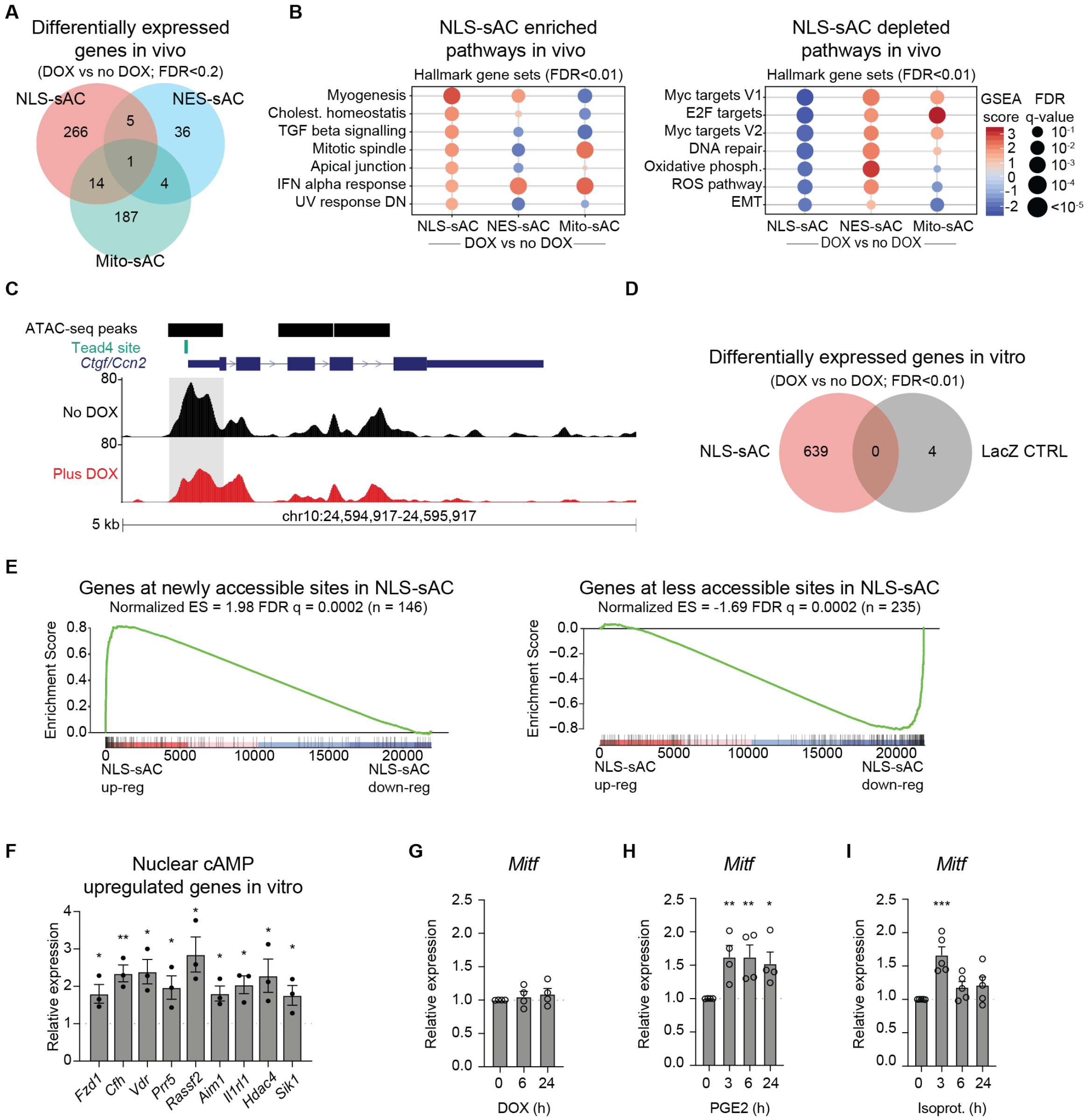


**Figure S5. Genomic analysis of cAMP microdomains.**
A) Venn diagram of differentially expressed genes in tumors formed following expression of NLS-, NES-, or Mito-sAC in melanoma cells by doxycycline (DOX), as compared to matched control (no DOX) for each microdomain.
B) GSEA of global expression changes in NLS-sAC (DOX) vs control (no DOX), NES-sAC (DOX) vs control (no DOX), or mito-sAC (DOX) vs control (no DOX) expressing tumors. Upregulated and downregulated panels represent gene sets with significant (FDR < 0.01) positive and negative enrichment, respectively, in NLS-sAC vs control. Extension of Figure 3A.
C) Example of ATAC-seq tracks showing reduction in accessibility of Tead4 binding site in the promotor region of *Ctgf/Ccn2* upon induction of nuclear cAMP with doxycycline (Plus DOX; 48h) as compared to control (no DOX).
D) Venn diagram of differentially expressed genes in NLS-sAC melanoma cell line and LacZ CTRL line after 48h doxycycline treatment in vitro.

E) GSEA enrichment plots for genes adjacent to differentially accessible DNA elements ranked by change in expression in NLS-sAC melanoma line after 48h doxycycline treatment. ES, enrichment score. FDR, false discovery rate.
F) RT-PCR confirmation of selected upregulated genes identified by RNA-seq on NLS-sAC melanoma line after 48h doxycycline treatment represented as DOX/no DOX mRNA level. Error bars, SEM; Student's t-test.
G) RT-PCR confirmation showing lack of *Mitf* gene induction by nuclear cAMP. n=4; error bars, SEM; ANOVA.
H) *Mitf* gene induction by tmAC-produced cAMP upon prostaglandin E2 (PGE2) treatment in mouse melanoma cell line. n=5; error bars, SEM; ANOVA.
I) *Mitf* induction by tmAC produced cAMP upon isoproterenol (Isoprot.) stimulation in mouse melanoma cell line. n=5; error bars, SEM; ANOVA.
(ns, $P > 0.05$; *, $P \leq 0.05$; **, $P \leq 0.01$; ****, $P \leq 0.0001$).

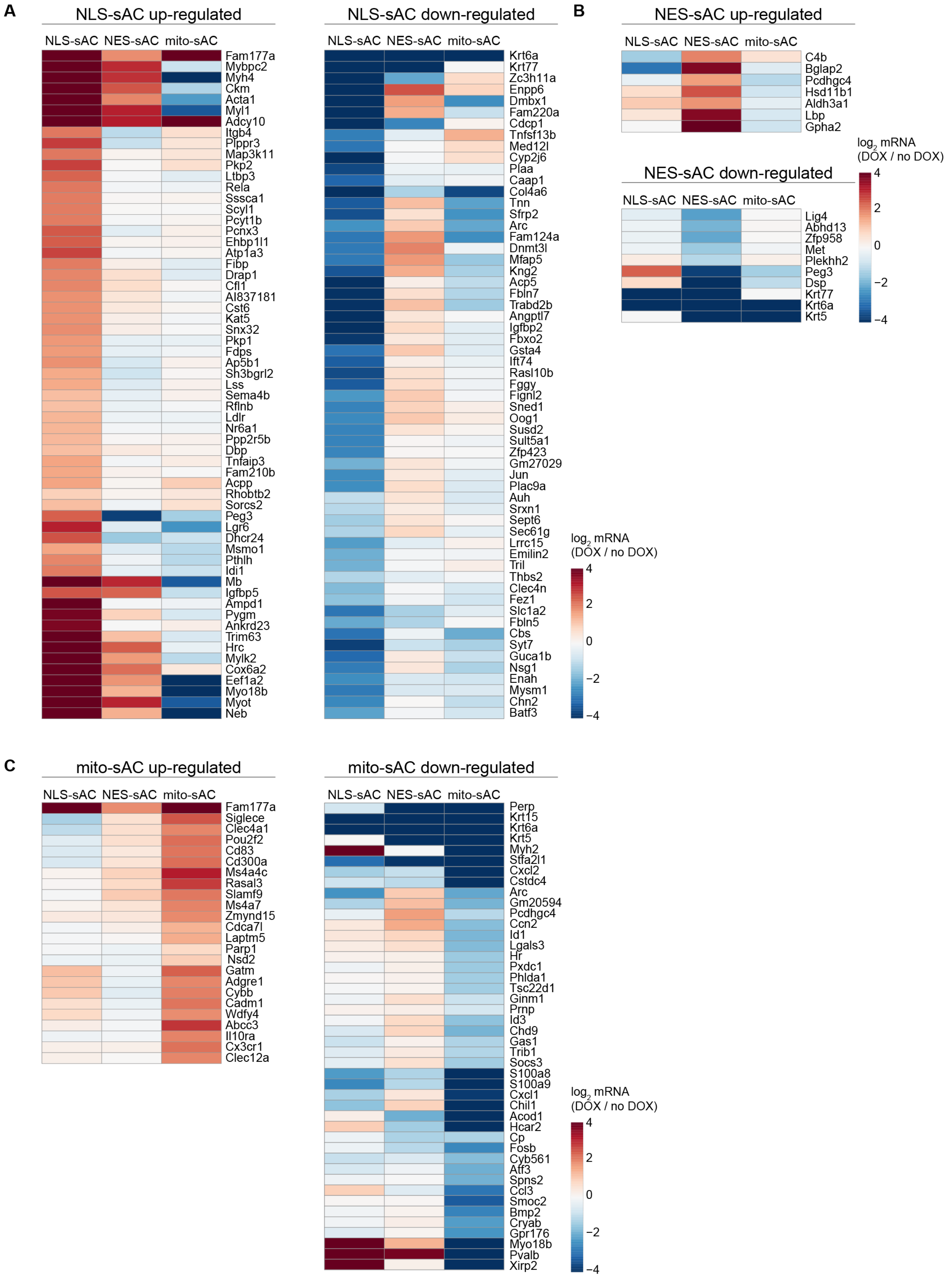
A
NLS-sAC up-regulated
NLS-sAC NES-sAC mito-sAC
Fam177a Mybpc2 Myh4 Ckm Acta1 Myl1 Adcy10 Itgb4 Plppr3 Map3k11 Pkp2 Ltbp3 Rela Sssca1 Scyl1 Pcyt1b Pcnx3 Ehbp1l1 Atp1a3 Fibp Drap1 Cfl1 AI837181 Cst6 Kat5 Snx32 Pkp1 Fdps Ap5b1 Sh3bgrl2 Lss Sema4b Rflnb Ldlr Nr6a1 Ppp2r5b Dbp Tnfaip3 Fam210b Acpp Rhobtb2 Sorcs2 Peg3 Lgr6 Dhcr24 Msmo1 Pthlh Idi1 Mb Igfbp5 Ampd1 Pygm Ankrd23 Trim63 Hrc Mylk2 Cox6a2 Eef1a2 Myo18b Myot Neb
NLS-sAC down-regulated
NLS-sAC NES-sAC mito-sAC
Krt6a Krt77 Zc3h11a Enpp6 Dmbx1 Fam220a Cdcp1 Tnfsf13b Med12l Cyp2j6 Plaa Caap1 Col4a6 Tnn Sfrp2 Arc Fam124a Dnmt3l Mfap5 Kng2 Acp5 Fbln7 Trabd2b Angptl7 Igfbp2 Fbxo2 Gsta4 Ift74 Rasl10b Fggy Fignl2 Sned1 Oog1 Susd2 Sult5a1 Zfp423 Gm27029 Jun Plac9a Auh Srxn1 Sept6 Sec61g Lrrc15 Emilin2 Tril Thbs2 Clec4n Fez1 Slc1a2 Fbln5 Cbs Syt7 Guca1b Nsg1 Enah Mysm1 Chn2 Batf3
log2 mRNA (DOX / no DOX)
4 2 0 −2 −4
B
NES-sAC up-regulated
NLS-sAC NES-sAC mito-sAC
C4b Bglap2 Pcdhgc4 Hsd11b1 Aldh3a1 Lbp Gpha2
log2 mRNA (DOX / no DOX)
4 2 0 −2 −4
NES-sAC down-regulated
NLS-sAC NES-sAC mito-sAC
Lig4 Abhd13 Zfp958 Met Plekhh2 Peg3 Dsp Krt77 Krt6a Krt5
C
mito-sAC up-regulated
NLS-sAC NES-sAC mito-sAC
Fam177a Siglece Clec4a1 Pou2f2 Cd83 Cd300a Ms4a4c Rasal3 Slamf9 Ms4a7 Zmynd15 Cdca7l Laptm5 Parp1 Nsd2 Gatm Adgre1 Cybb Cadm1 Wdfy4 Abcc3 Il10ra Cx3cr1 Clec12a
mito-sAC down-regulated
NLS-sAC NES-sAC mito-sAC
Perp Krt15 Krt6a Krt5 Myh2 Stfa2l1 Cxcl2 Cstdc4 Arc Gm20594 Pcdhgc4 Ccn2 Id1 Lgals3 Hr Pxdc1 Phlda1 Tsc22d1 Ginm1 Prnp Id3 Chd9 Gas1 Trib1 Socs3 S100a8 S100a9 Cxcl1 Chil1 Acod1 Hcar2 Cp Fosb Cyb561 Aff3 Spns2 Ccl3 Smoc2 Bmp2 Cryab Gpr176 Myo18b Pvalb Xirp2
log2 mRNA (DOX / no DOX)
4 2 0 −2 −4

**Figure S6. Differential impact of cAMP microdomains on gene expression.**
Heat maps showing the $Log_2$ fold change for the most significant upregulated and downregulated genes in NLS-sAC (DOX vs no DOX) (A), NES-sAC (DOX vs no DOX) (B), and mito-sAC (DOX vs no DOX) (C). Differentially expressed genes (FDR < 0.05) specific to each microdomain are shown. Gene expression in the other respective microdomains in each figure panel is shown for comparison.

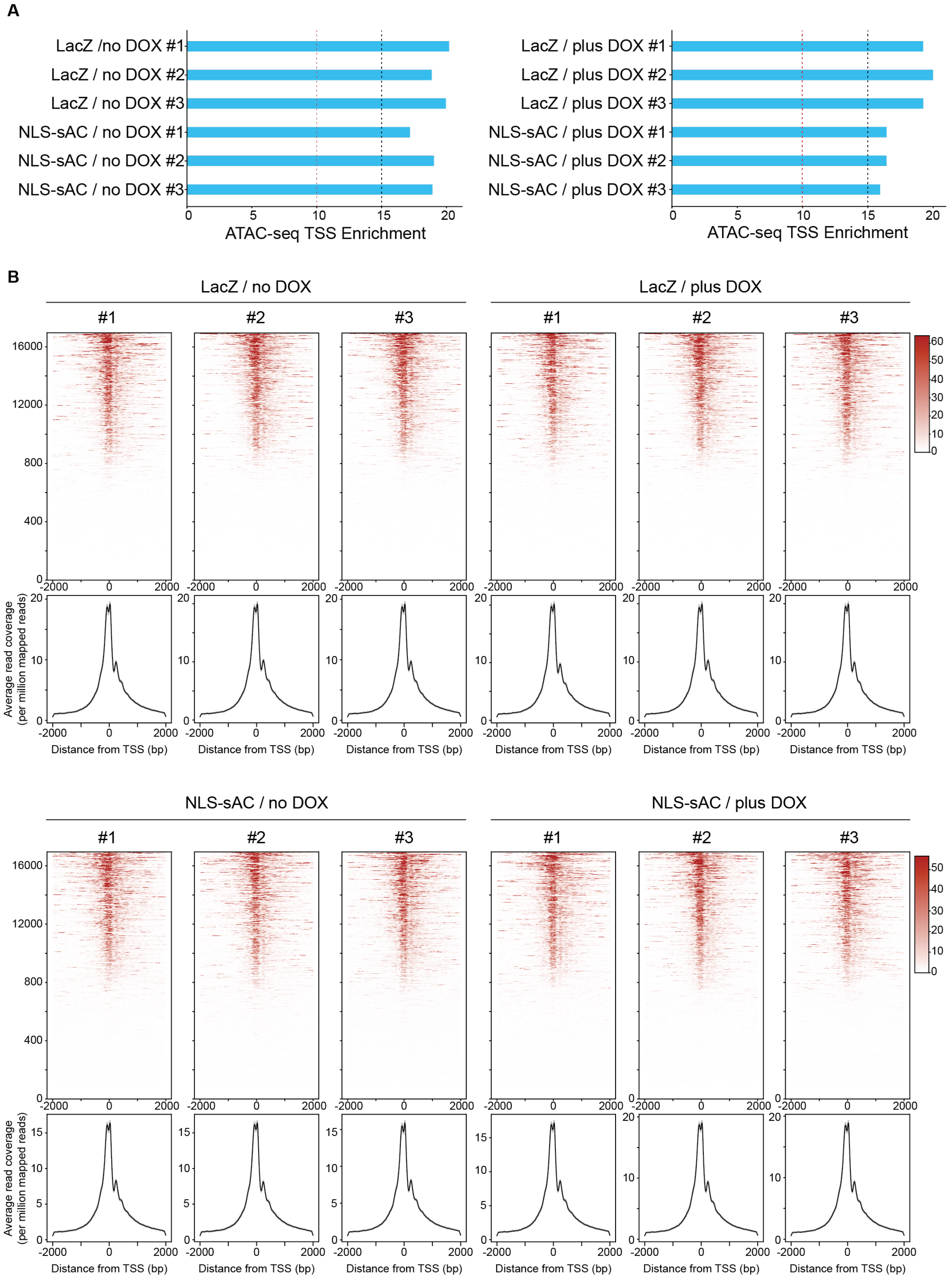
A
LacZ /no DOX #1
LacZ / no DOX #2
LacZ / no DOX #3
NLS-sAC / no DOX #1
NLS-sAC / no DOX #2
NLS-sAC / no DOX #3
ATAC-seq TSS Enrichment
LacZ / plus DOX #1
LacZ / plus DOX #2
LacZ / plus DOX #3
NLS-sAC / plus DOX #1
NLS-sAC / plus DOX #2
NLS-sAC / plus DOX #3
ATAC-seq TSS Enrichment
B
LacZ / no DOX
LacZ / plus DOX
NLS-sAC / no DOX
NLS-sAC / plus DOX
#1
#2
#3
Average read coverage (per million mapped reads)
Distance from TSS (bp)

**Figure S7. Quality control of ATAQ-seq.**
A) ATAC-seq TSS enrichment scores for each sample. Vertical lines indicate the ENCODE quality guidelines thresholds (>10 acceptable; >15 ideal).
B) Heatmaps of the ATAC-seq read densities with corresponding metagene plots for each sample (biological replicates +/- doxycycline are indicated above each heatmap).

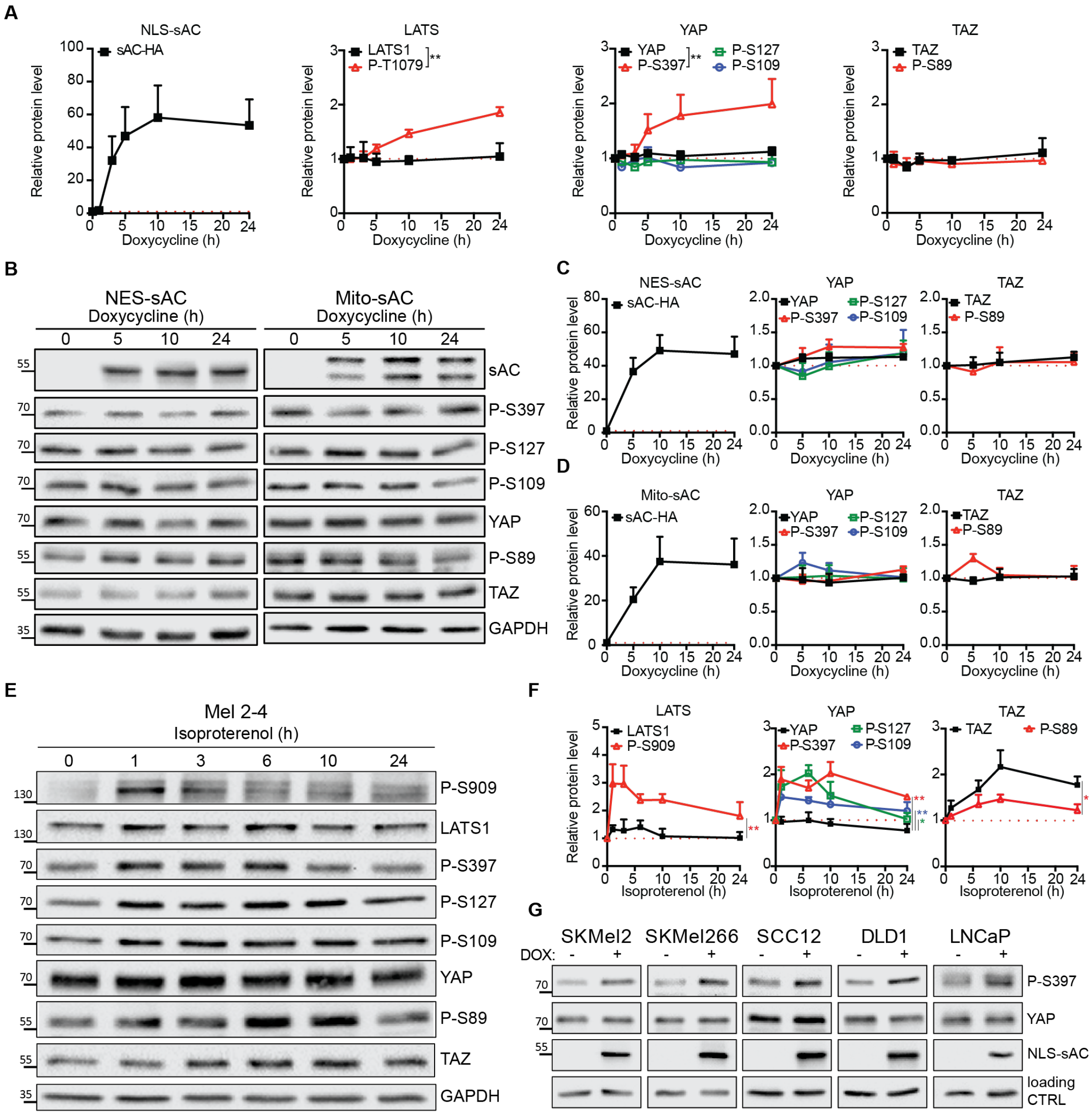


**Figure S8. Key regulators of Hippo pathway are differently affected by distinct cAMP microdomains.**

A) Quantification of immunoblot signal following NLS-sAC induction related to Figure 4B. Error bars, SEM; Mixed-effect ANOVA; n≥3.

B) Western blot analysis showing lack of YAP or TAZ phosphorylation in response to activation of either cytoplasmic (NES-sAC) or mitochondrial (Mito-sAC) cAMP microdomains.

C) Quantification of immunoblot signal related to Figure S8B (left panel, NES-sAC). Error bars, SEM; ANOVA; n≥3.

D) Quantification of immunoblot signal related to Figure S8B (right panel, Mito-sAC). Error bars, SEM; ANOVA; n≥3.

E) Western blot analysis showing that tmAC activation by the GPCR agonist isoproterenol leads to phosphorylation of LATS1, YAP, and TAZ at multiple residues.

F) Quantification of immunoblot signal related to Figure S8E. Error bars, SEM; ANOVA; n≥3.

G) Western blot analysis of YAP phosphorylation at S397 following induction of nuclear cAMP in a panel of human cancer cell lines. DOX, doxycycline. Loading CTRL in SKMel2, SKMel266, SCC12, and DLD1 panels is GAPDH, and in LNCaP is Beta-Tubulin.

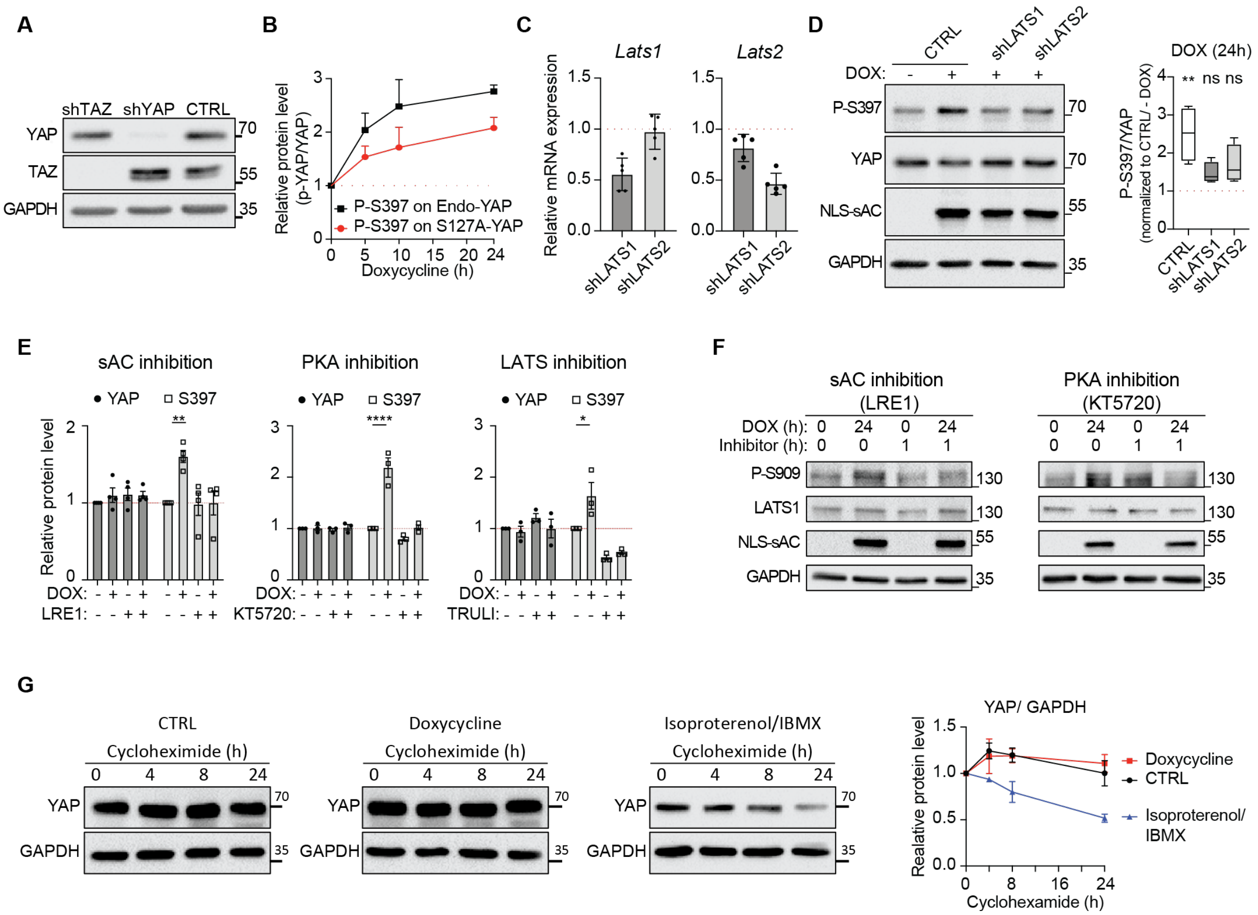

**Figure S9. YAP but not TAZ is regulated by nuclear cAMP microdomain through LATS1/2 phosphorylation.**

A) Western blot analysis of YAP and TAZ following shRNA knock down.
B) Quantification of immunoblot signal related to Fig. 4E. n=3; P-S397 was normalized to the total YAP level, either endogenous wild type YAP (Endo-YAP), or overexpressed S127A-YAP mutant.
C) RT-PCR confirmation of *Lats1* or *Lats2* knock down by shRNA.
D) Western blot analysis of S397 phosphorylation upon nuclear cAMP induction in melanoma cells with either LATS1 or LATS2 knock-down (shLATS1 and shLATS2, respectively). Right panel, quantification of Western blot band volume of P397 YAP ratio compared to total YAP normalized to control media (-, no DOX). Combination of 4 independent experiments; boxes represent 25-75 percentile, while whiskers show min. to max. values; the line within boxes indicates median.
E) Quantification of immunoblot signal relative to control conditions (-,-) related to Fig. 4G. sAC inhibition, n=4; PKA inhibition, n=3; LATS inhibition, n=3. DOX, doxycycline; LRE1, sAC inhibitor, 50 µM; KT5720, PKA inhibitor, 10 µM; TRULI, LATS inhibitor, 500 nM. Error bars, SEM; Student's t-test.
F) Western blot analysis examining phosphorylation status of LATS1 kinase in cells treated with either 50 µM LRE1 (sAC inhibitor) or 10 µM KT5720 (PKA inhibitor) in the presence or absence of doxycycline (DOX).

G) Western blot analysis of YAP in mouse melanoma cells in the presence of cycloheximide over time in either untreated (CTRL), pretreated with 1 μg/ml doxycycline (Doxycycline) for 24h to induce the nuclear cAMP microdomain, or treated with 10 μM isoproterenol and 500 μM IBMX (Isoproterenol/IBMX) for 24h to induce tmACs. Right panel shows change in YAP protein level, relative to GAPDH and normalized to timepoint 0 (n=5 for CTRL and Doxycycline; n=3 for Isoproterenol/IBMX).
(*, $P \leq 0.05$; **, $P \leq 0.01$; ****, $P \leq 0.0001$).

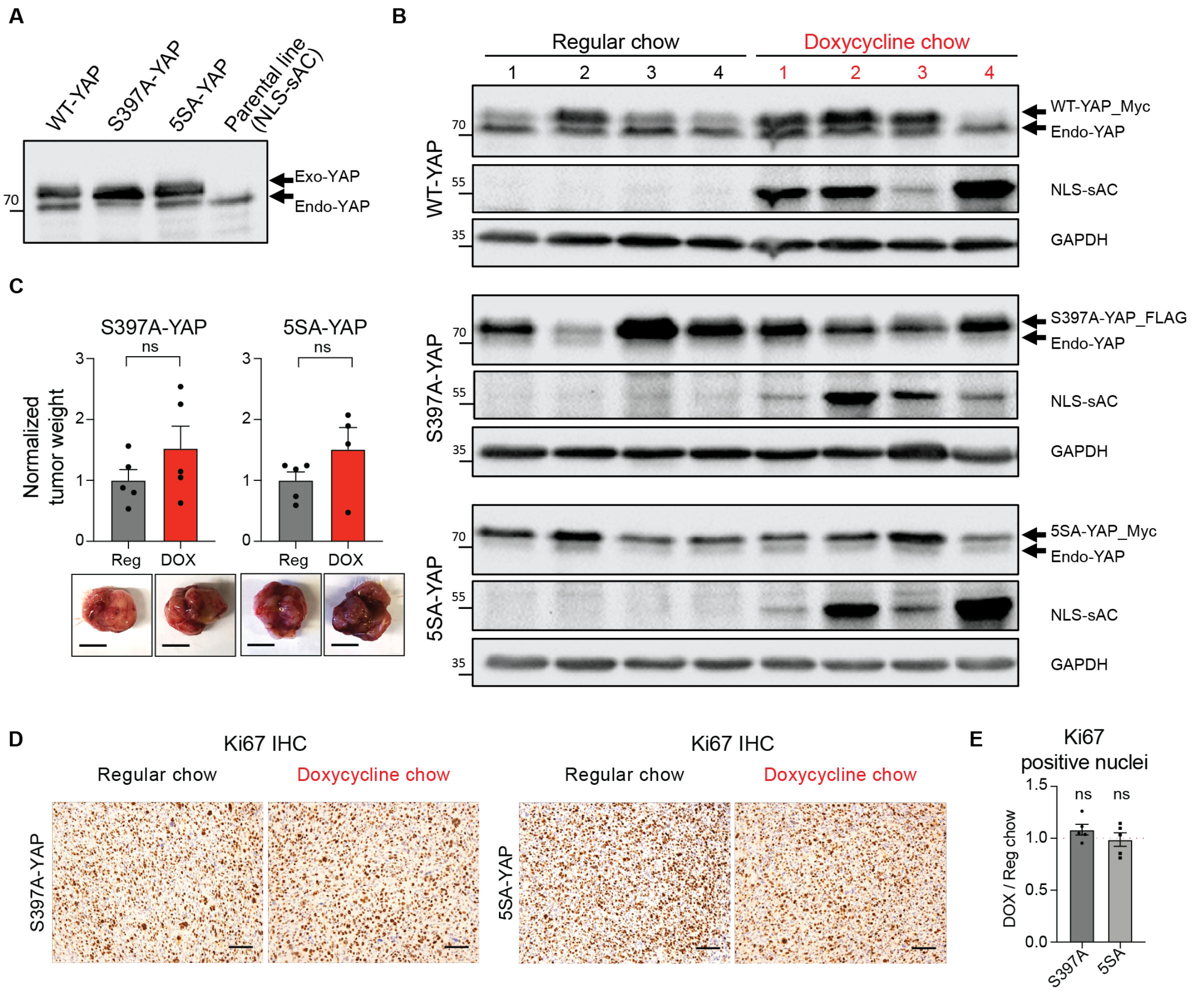


**Figure S10. YAP S397 phosphorylation and tumor suppression by nuclear cAMP.**
A) Confirmation of YAP overexpression in NLS-sAC cell line after transduction with viruses encoding YAP variants: WT-YAP (wild type), S397A-YAP, and 5SA-YAP (five key serines mutated to alanine). Western blot detection with anti-YAP antibody that binds to both endogenous (endo) and overexpressed exogenous (exo) YAP variant as indicated by arrows.
B) Western blot analysis of tumor lysates, confirming NLS-sAC induction by doxycycline chow and expression of exogenous YAP isoforms.
C) Bar graphs showing normalized tumor weight with corresponding gross image examples from the experiment in Fig. 5B. Mean shown; Error bars, SEM; Student's t-test; scale bar 1 cm.
D) Microscopic image examples of Ki67 immunohistochemistry (IHC) analysis of tumor sections. Scale bar 100 µm.
E) Quantitation of Ki67 positive nuclei in tumor sections represented as a fold change doxycycline chow (DOX) over regular chow (REG) cohort. Mean values with data points for individual tumors shown. Error bars, SEM; Student's t-test; ns, P > 0.05.
